# Automated and parallelized spike collision tests to identify spike signal projections

**DOI:** 10.1101/2021.10.21.465238

**Authors:** Keita Mitani, Masanori Kawabata, Yoshikazu Isomura, Yutaka Sakai

**Affiliations:** Brain Science Institute, Tamagawa University, Machida, Tokyo, Japan; Department of Physiology and Cell Biology, Graduate School of Medical and Dental Sciences, Tokyo Medical and Dental University, Tokyo, Japan

## Abstract

The spike collision test is a highly reliable technique to identify axonal projection of a neuron recorded electrophysiologically for investigating functional spike information among brain areas. It is potentially applicable to more neuronal projections by combining multi-channel recording with optogenetic stimulation. Yet, it remains inefficient and laborious because an experimenter must visually select spikes in every channel and manually repeat spike collision tests for each neuron serially. Here, we established a novel technique to automatically perform spike collision tests for all channels in parallel (*Multi-Linc* analysis), employing two distinct protocols implemented in a multi-channel real-time processing system. The rat cortical neurons identified with this technique displayed physiological spike features consistent with excitatory projection neurons. Their antidromic spikes were similar in shape but slightly larger in amplitude compared with spontaneous spikes. In addition, we demonstrated simultaneous identification of reciprocal or bifurcating projections among cortical areas. Thus, our *Multi-Linc* analysis will be a powerful research approach to elucidate interareal spike communication.

---

The brain performs various functions by conveying spike signals of individual neurons cooperatively among brain areas. To elucidate such interareal spike communication, it is essential to examine spike activity of a projection neuron that is proved to send its axon to specific target areas. Extracellular spike (unit) recording is currently the only method that precisely captures every spike in any brain area of a behaving animal. The spike collision test can reliably determine the axonal projection of an extracellularly recorded neuron without requiring visualization (Bishop et al., 1962; Lipski, 1981). This test is based on the principle that two spikes always disappear if they collide with each other on the same axon. When we stimulate the axon in the target area, an evoked “antidromic” spike will be detected at the soma with a delay (**Fig. 1a**, upper; typically a few to tens of milliseconds later). Then, if the antidromic spike is evoked immediately after a spontaneous (“trigger”) spike is detected at the soma, they will collide and disappear on the axon and the antidromic spike will not be detected (**Fig. 1a**, middle). In contrast, if the recorded neuron is excited by other activated neurons via synapses, an evoked “synaptic” spike will be detected despite the trigger spike (**Fig. 1a**, lower). On the basis of the “success” of spike collision, the target area of the extracellularly recorded neuron can be identified. For the past 50 years, this analysis has been used to identify projection neurons in various brain areas, leading to fruitful findings on neural circuitry (e.g., Evart, 1968; Wilson, 1987; Turner & DeLong, 2000; Everling & Munoz, 2000). However, this classical analysis is extremely difficult and inefficient for technical reasons. First, it is laborious to search a single neuron with evoked spikes by advancing a recording electrode little by little. Second, electrical stimulation generates a huge electrical artifact that obstructs the spike detection. Third, the stimulation is prone to make an electrolytic lesion or insensitivity near the tip of the electrode.

**Fig. 1.**
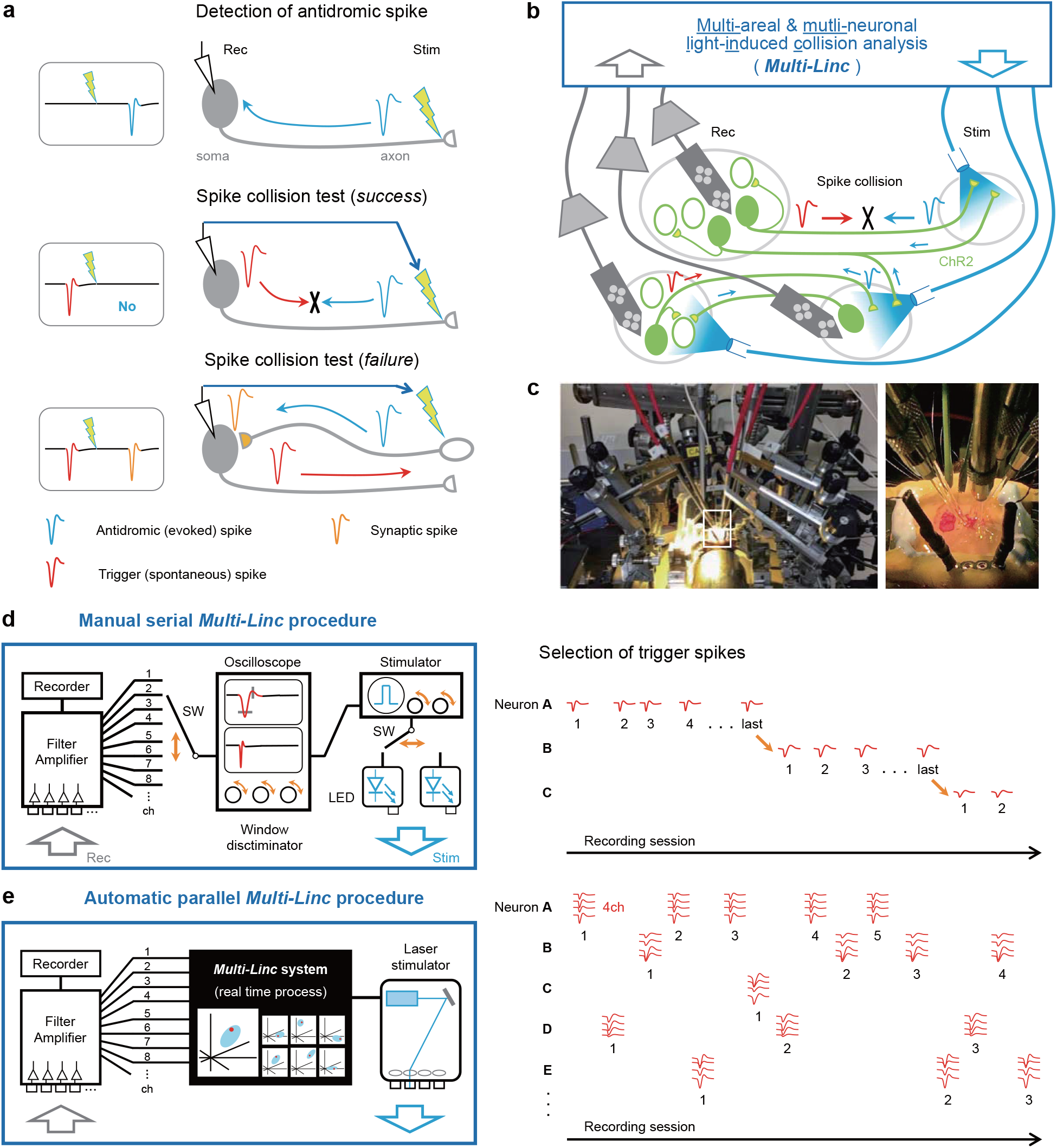
Automation and parallelization of the spike collision test. **a**, Principle of the spike collision test. An evoked antidromic spike (upper) disappears upon colliding with a spontaneous (trigger) spike on the same axon (middle). Otherwise, a synaptic spike persists (lower). **b**, The concept of the *Multi-Linc* analysis. Multi-unit recordings (Rec) are combined with optogenetic stimulation (Stim) to test spike collisions for multiple projection neurons among brain areas. **c**, A *Multi-Linc* experiment in a rat (left). Several optical fibers and multi-channel silicon probes were placed in different cortical areas (right). **d**, Schematic of manual and serial *Multi-Linc* procedures (left). An experimenter needs to continually operate analog-type devices visually and manually (orange arrows; input channel selector, window discriminator, stimulator, LED selector, etc.) to detect antidromic-like spikes and test spike collisions during the experiment. Consequently, neurons are subject to spike detection and spike collision tests one-by-one serially (right). See Saiki et al. (2018). **e**, Schematic of the automatic and parallel *Multi-Linc* procedure (left). A multi-channel *Multi-Linc* system automatically conducts real-time processes to isolate trigger spikes precisely using multi-dimensional spike features (every tetrode from as many as 128 channel inputs) and to select an optimal stimulus to test spike collision within several milliseconds. Many neurons can be efficiently subjected to spike collision tests in parallel (right).

Recently, several laboratories have employed multi-channel recording and optogenetic stimulation for the spike collision test to solve the aforementioned problems (Jennings et al., 2013; Li et al., 2015; Economo et al., 2018; Saiki et al., 2018). For example, we proposed a concept of “<u>multi-</u>areal and <u>multi-</u> neuronal light-<u>in</u>duced <u>c</u>ollision” (*Multi-Linc*) analysis, which repeats to test spike collisions for axonal projections of multiple neurons from multiple areas to other areas all at once (**Fig. 1b**; Saiki et al., 2018). To prioritize the efficiency of *Multi-Linc* analysis, as many spike collisions as possible are tested even though they are tentative and inaccurate *online* at the experimental stage; they are later re-evaluated more accurately to judge success *offline* at the analytical stage. This analysis has been used to identify distinct types of cortical (Soma et al., 2017; Saiki et al., 2018; Rios et al., 2019; Hamada et al., 2021) and striatal (Nonomura et al., 2018) projection neurons by combining multi-unit recording through tetrodes with optogenetic stimulation in behaving rats expressing Channelrhodopsin-2 (ChR2) (**Fig. 1c**).

In the original *Multi-Linc* procedure, an experimenter selects one recording channel and one stimulation site manually and searches the trigger spike matched with one of the evoked spikes on an oscilloscope visually and manually (**Fig. 1d**, left, orange arrows; Saiki et al., 2018). Consequently, the experimenter must wait for tens of spike collision trials to complete in a single neuron before searching the next neuron (**Fig. 1d**, right). Such manual series operations and visual spike isolation seriously degrade the efficiency and accuracy, respectively, of multi-channel spike collision tests. If trigger spikes are precisely selected in a multi-dimensional space of spike features for each tetrode automatically within several milliseconds (**Fig. 1e**, left), spike collision tests can be conducted in parallel for multiple neurons (F**ig. 1e**, right). Thus, the automation and parallelization of the *Multi-Linc* procedure are expected to improve the efficiency and accuracy of spike collision experiments. The key technical issues to be solved are 1) automatic detection of inferred antidromic spikes from noisy background signals, 2) real-time selection of trigger spikes for spike collision tests, and 3) implementation of reliable real-time processing (on a millisecond time scale) in a computer system with multi-channel inputs and outputs.

In the present work, we have overcome these technical issues in both hardware and software and have established an automatic and parallel *Multi-Linc* system that is real-time computerized for multi-channel spike collision tests. Using this novel system, we succeeded in automatically identifying cortical projection neurons recorded in the motor cortex of ChR2-expressing transgenic rats. As far as we know, this study is the first to report the automation and parallelization of spike collision tests for multiple neurons. We expect our *Multi-Linc* analysis method to provide insights into the principle of fast spike communication among brain areas in the future.

## Results

We composed the automatic and parallel *Multi-Linc* procedure of three serial steps for online control of experiments (**Fig. 2a**; see Online Methods for details): collection of evoked responses, inference of antidromic spikes in evoked responses, and spike collision test for inferred antidromic spikes. The first two steps correspond to the antidromic spike detection in **Fig. 1a**, top, and the third corresponds to the spike collision test in Fig. 1a, middle and bottom. In the evoked response collection (**Fig. 2a**, first block), one of the candidate projection sites in which optical fibers are placed is selected and stimulated optogenetically at an arbitrary timing. Simultaneous recordings through multi-tetrode probes inserted into multiple brain areas (Saiki et al., 2018; Kawabata et al., 2020) are stored for several tens of milliseconds after the stimulation (evoked responses). The slow component of the change in potential is removed by a high-pass filter such that only the spike activity remains. The next stimulation site and timing are selected after a certain interval (∼1 sec) to avoid residual effects of artificially evoked neural activity. The evoked response collection (first block in **Fig. 2a**) is completed after a sufficient number of stimulations per stimulation site (typically 100 times per site). The set of evoked responses is then obtained for every combination of tetrodes and stimulation sites, which is delivered to the next step (second block in **Fig. 2a**).

**Fig. 2.**
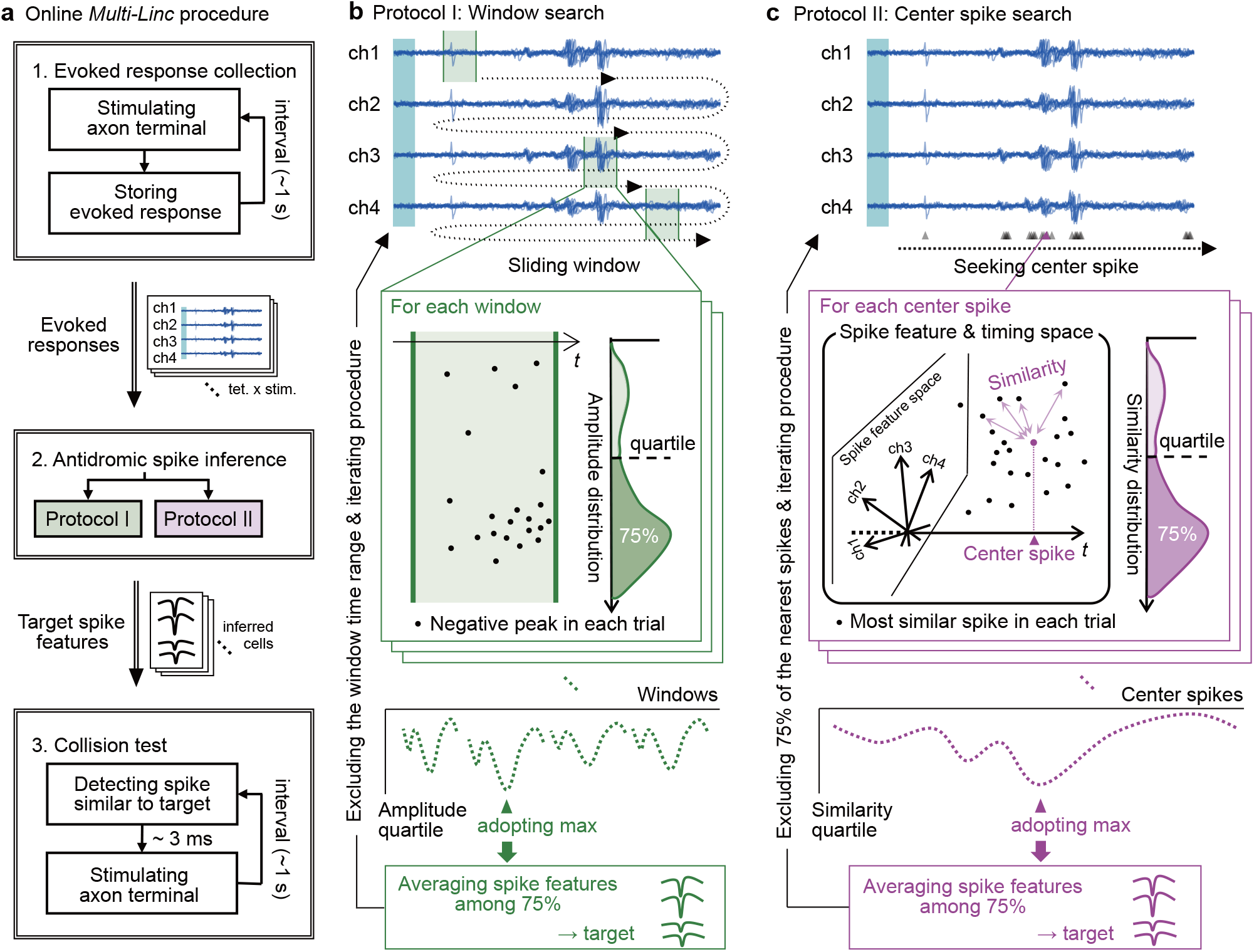
Automatic and parallel *Multi-Linc* procedure. **a**, Flow of the online *Multi-Linc* procedure. **b, c**, Two protocols to infer antidromic spikes of an identical neuron among multiple evoked spikes. The schema corresponds to a procedure on a certain combination of a tetrode and a stimulation site. **b**, Window search protocol (protocol I). Evaluation of evoked negative peaks within a sliding time window determines candidate antidromic spikes. **c**, Center spike search protocol (protocol II). All evoked spikes in every stimulation trial are sought. The similarity of each spike to the most similar spike in every other trial is evaluated. “Similarity” is defined in a space of spike features and timing. Typically, the peak pattern of a spike in four channels of a tetrode can form a space of spike features. Evaluation of aggregation in a high-dimensional space of the evoked spike feature and timing determines the candidate antidromic spikes.

If a neuron around a tetrode projects to a stimulation site, antidromic spikes might be observed in the evoked responses. However, the evoked responses typically contain many synaptic spikes through the neuronal population. Occasionally, antidromic spikes of different neurons might be mixed. The purpose of the antidromic spike inference (second block in **Fig. 2a**) is to sort a group of spikes likely to be antidromic spikes of an identical neuron from contaminated multiple spikes in the noisy evoked responses. If synaptic spikes or antidromic spikes of other neurons contaminate the inferred group of spikes, then incorrect spontaneous spikes are likely to be selected as triggers for spike collision tests, which would decrease the success rate of spike collisions. Therefore, the accuracy of antidromic spike inference is extremely important in the online *Multi-Linc* procedure. Antidromic spikes should be observed with stable latency from stimulation timings because the spike conduction time should be determined by the excitability and length of the axon. By contrast, synaptic spikes might be evoked at various timings depending on the states of the neuronal population. The problem lies in the difficulty associated with segregating temporally aggregated spikes from noisy timing spikes. To solve this problem, we adopted two types of protocols: “window search” (protocol I; **Fig. 2a**, second block, green) and “center spike search” (protocol II; magenta). These two protocols were also used in offline judgement of spike collision after experiments (**Supplementary Fig. 1a**).

In protocol I, temporal aggregation is evaluated using a short temporal window independently for the four channels of a tetrode. For each combination of a tetrode and a stimulation site, the temporal window is slid along time in the filtered evoked responses from channels 1 to 4 (**Fig. 2b**, top). A spike waveform exhibits a sharp negative peak in extracellular recording. Thus, stably large negative peaks within a short temporal window indicate temporally aggregated spikes. For each window, negative peaks (dots in **Fig. 2b**, middle left) are determined in respective trials of stimulations and the distribution of peak amplitudes is obtained (**Fig. 2b**, middle right). The goodness of the window can be characterized with the first quartile of the amplitude (75% from the largest) in the amplitude distribution because a larger amplitude quartile indicates a more stable occurrence of spikes within the window. All the windows of timings and channels are characterized in the same manner. The windows with amplitude quartiles below a certain threshold are excluded from candidates. Among the remaining candidates, the window with the maximum amplitude quartile is adopted as the best window (**Fig. 2b**, bottom). Here, we used the negative peaks with an amplitude greater than the amplitude quartile (75% from the largest) within the adopted window as typical spike waveforms likely to be antidromic spikes of an identical neuron. The spike waveforms corresponding to the 75% of negative peaks are averaged for the representative waveform of inferred antidromic spikes. Although the adopted window is defined in a certain channel of the tetrode, the waveform patterns for all four channels are averaged. The feature of the averaged waveform pattern is then delivered to the next step (**Fig. 2a**, third block) as the target of the spike collision test. The windows overlapping the time range of the adopted window are excluded from candidates; the selection of the best window among the remaining candidates is then iterated.

In protocol II, not only temporal aggregation but also aggregation of spike waveform features is evaluated on the basis of the similarity between spikes. Spike timings are detected within the evoked responses for all trials of stimulations (triangles in **Fig. 2c**, top). On the basis of timings and waveform features of the detected spikes, aggregation of spikes is searched, and the center spike of the aggregation is determined. For each spike, the nearest spikes in respective trials are determined (dots in **Fig. 2c**, middle). The similarity between spikes is defined in the multi-dimensional space of spike timing and features (see Online Methods). The degree of aggregation can be characterized by the first quartile in the similarity distribution (75% from the most similar one; **Fig. 2c**, middle right). All detected spikes are characterized with the similarity quartiles in the same manner. Spikes with the similarity quartiles below a certain threshold are excluded from candidates for the center spike. Among the remaining candidates, the center spike with the maximum similarity quartile is adopted as the center of the aggregation (**Fig. 2c**, bottom). In the same manner as protocol I, protocol II uses 75% of the spikes nearest the adopted center spike as typical spike waveforms likely to be antidromic spikes of an identical neuron. The waveform patterns of the 75% are averaged for the target of the spike collision test. Once adopted, the 75% of the nearest spikes are excluded from candidates, and selection of the center spike is iterated.

In the spike collision test (**Fig. 2a**, third block), spontaneous spikes are detected in real time and in parallel on all tetrodes. If a detected spontaneous spike is close to either of the target spike features in the same tetrode, then the spike is adopted as a trigger for the collision test. The stimulation site attributed to the target is immediately stimulated. The stimulation should start within a few milliseconds after the occurrence of the trigger spike for the spike collision to be successful. This process requires the highest performance of real-time processing. Acquiring data from multiple tetrodes, filtering all channels, detecting spikes, determining the similarities to target spikes, and exciting a laser require time. We implemented algorithms for easy filtering and spike detection and selected hardware components to achieve stimulation within 3.2 ms, acquiring data from 32 tetrodes (128 channels).

We conducted experiments using the online *Multi-Linc* controller (**Fig. 2a**) by implementing either protocol I or II in the bilateral motor cortices of ChR2-expressing transgenic rats. Both protocols worked well and we collected a number of sessions of data (**Supplementary Table 1**). After the experiments, we inferred antidromic spikes in the whole stimulation trials offline using both protocols I and II, irrespective of the protocol used in the online controller (**Supplementary Figs. 1 and 2**; see Online Methods). We also detected and sorted spikes on each tetrode into clusters of putative identical neurons throughout the experimental session by using a standard method in multi-unit recordings (Takekawa et al., 2012). We judged whether each neuron cluster might pass the collision tests for the inferred antidromic spikes and found neurons such that the projection targets were successfully identified (**Fig. 3a**). As a result, multiple projections were simultaneously identified in numerous sessions by both online controllers I and II (left and right in **Fig. 3b**, respectively). The numbers of projections successfully identified in offline judgements I and II were equivalent (green and magenta in **Fig. 3b**, respectively). The majority succeeded in both judgements I and II (**Supplementary Fig. 3a**). Thus, the results were assured by different criteria. If we also accept a projection judged as success in only one of protocols I and II, then the successful projections exceeded ten per session at maximum in both online controller I and II (**Supplementary Fig. 3b**). We showed the evoked traces with antidromic spikes and with spike collisions for all 13 identified projections from 12 neurons (two different projections were identified in one of the neurons) in the maximum example obtained by online controller I (**Supplementary Fig. 4**). These were results of the first use of the online *Multi-Linc* controller; the control parameters have not been fine tuned. Hence, the maximum outcomes showed that fine tuning on the basis of the properties of successful neurons should enable the stable identification of more than ten neurons per session. Hereafter, we explain the details of the judgements and the properties of the data sessions collected with online controllers I and II.

**Fig. 3.**
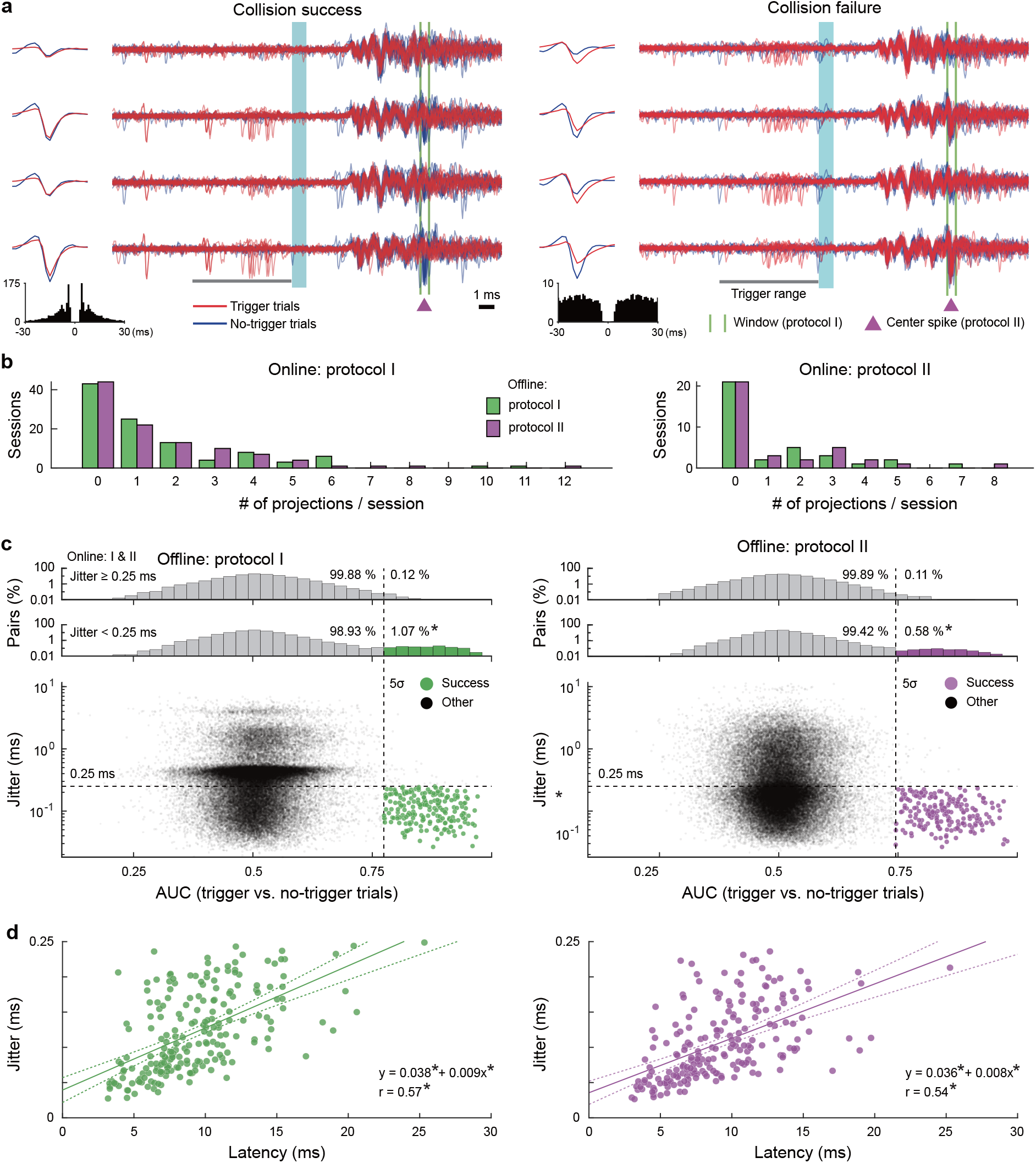
Offline judgement for identification of neuronal projections. **a**, Examples of successful (left) and unsuccessful (right) pairs for collision tests, which correspond to different example neurons (spike clusters) tested for a common antidromic spike inference by protocol I (the window indicated by green vertical lines) and protocol II (the center spike indicated by a magenta triangle). Traces recorded by a four-channel tetrode around stimulation timings (light blue) in trigger (red) and no-trigger (blue) trials were overlayed. Median waveforms of trigger spontaneous spikes (red) and evoked spikes in no-trigger trials (blue) are plotted on the left sides. The autocorrelogram of spontaneous spikes was shown in the bottom of each. Spontaneous spikes of the successful neuron (left) within the trigger range (gray bar) eliminated inferred antidromic spikes to be evoked (within the green lines, or around the magenta triangle), whereas spikes of the unsuccessful neuron (right) could not eliminate the antidromic spikes. **b**, The number of successfully identified neuronal projections per session (left, online controller by protocol I; right, online controller by protocol II). The outcomes of offline judgements by protocols I and II are plotted as the green and magenta bars, respectively. **c**, The definitions of offline judgements by protocols I (left) and II (right). The data sessions obtained by online controllers I and II were gathered. Each dot in the scatter plots indicates the area under the receiver operating characteristic curve (AUC) between trigger and no-trigger trials, which was used as a variable to quantify the elimination of evoked spikes and the jitter of evoked spikes in no-trigger trials (the quartile deviation of the latency) in each tested pair. Upper histograms of AUCs are shown in logarithmic scale separately for ones with jitter greater than and less than 0.25 ms (horizontal dashed lines; asterisks (*) indicate *p* < 0.005 in 2×2 χ^2^ tests). Green and magenta dots indicate tested pairs judged to pass collision tests on the basis of the defined criteria (AUC > 5σ and jitter < 0.25 ms), and gray dots indicate the other pairs. **d**, The distribution of jitters and latencies of antidromic spikes for the identified neuronal projections. The regression lines (solid) and the 95% confidence intervals (dashed curves) are shown. Asterisks (*) mean *p* < 0.005 in Pearson’s correlation tests and linear regression tests.

We adopted broad inferences of antidromic spikes, including ones with relatively long jitters, for control comparisons. A set of antidromic spikes inferred to originate in an identical neuron was a target of identification. If the inferred antidromic spikes of an identification target are true antidromic spikes of a certain neuron, then the jitter should be relatively short and a spontaneous spike of the neuron immediately before stimulation should eliminate the evoked antidromic spike by collision in the axon (e.g., **Fig. 3a**, left). We searched such pairs of neurons and identification targets among all the pairs. For each pair, we extracted the trigger stimulation trials in which spontaneous spikes were observed in the time range to cause collision (red traces in **Fig. 3a**; see **Supplementary Fig. 1b** and Online Methods). If the number of extracted trigger trials was less than 15, then the pair was excluded for analyses. We also extracted no-trigger trials without spikes in the time range to enable collision for control comparison (blue traces in **Fig. 3a**; see **Supplementary Fig. 1b**). To ensure a fair comparison, we extracted the no-trigger trials only from trials temporally close to the respective trigger trials in a recording time course. We judged the spike collision on the basis of discrimination between the trigger and no-trigger trials. We also quantified the jitter of evoked spike timings with the quartile deviation among the extracted no-trigger trials. The jitter of synaptically evoked spikes is known to be longer than antidromic ones. On the basis of previous studies on antidromic stimulations by optogenetics (Li et al., 2015; Saiki et al., 2018), we regarded the jitters longer than 0.25 ms (above the horizontal dashed lines in **Fig. 3c**) as controls that could feasibly be synaptic ones.

The elimination of an antidromic spike to be evoked in each trial can be judged by the variables used for the antidromic spike inference—specifically, the peak amplitude within the inferred window in protocol I and the similarity to the inferred center spike in protocol II (see **Fig. 2b,c** and **Supplementary Figs. 1c,d and 2e,f**). We applied the receiver operating characteristic (ROC) analyses for the amplitude or the similarity to quantify discrimination between trigger and no-trigger trials. Because the majority of the tested pairs of neurons and identification targets should not match, the discrimination would be non-significant in the majority of pairs. Actually, the areas under the curves (AUCs) of the ROC were distributed around 0.5, the chance level (horizontal values of gray dots in **Fig. 3c**). For jitters shorter than 0.25 ms (horizontal dashed lines), however, the AUC distribution exhibited a long right tail (upper histograms). There were significantly more outliers of the AUC greater than five times the standard deviation (AUC > 5σ) for the shorter jitters (<0.25 ms) compared with the fraction for longer jitters (>0.25 ms; *p* = 6.5 × 10^−57^ in protocol I, *p* = 1.9 × 10^−18^ in protocol II, χ^2^ test). These outliers are likely to be pairs such that the spontaneous spikes of the neurons might eliminate the antidromic spikes to be evoked. We then adopted the pair to be succeeded in the collision test such that the AUC was greater than 5σ and the jitter was shorter than 0.25 ms (magenta and green dots in **Fig. 3c**). We confirmed the validity of our judgement in the original scale of the peak amplitude and the similarity (**Supplementary Fig. 5**).

The latencies of successfully identified antidromic spikes were distributed around 10 ms (**Fig. 3d**). The jitters were linearly correlated with the latencies (*r* = 0.57, *p* = 1.2 × 10^−44^ in protocol I; *r* = 0.54, *p* = 5.1 × 10^−33^ in protocol II, Pearson’s correlation test). If it is necessary to identify projections with latencies longer than 20 ms, then the criterion for jitters (<0.25 ms) could be weakened. Actually, we could still observe AUC outliers at >0.25 ms jitter (**Fig. 3c**). Here, we adopted a strict criterion to ensure validity; however, an appropriate jitter criterion that depends on the latency may enable more efficient identification.

We also checked the cell types of successful neurons on the basis of the waveforms of spontaneous spikes. Fast-spiking neurons—a subtype of cortical neurons that exhibit narrow spike waveforms—are interneurons known to not project on other brain areas (Isomura et al., 2009). Actually, we observed a bimodal distribution of trough-to-peak durations in average spike waveforms of unsuccessful (other) neurons (gray dots and histograms in **Fig. 4a**). The narrower group exhibited a higher ongoing spike rate, consistent with the properties of fast-spiking neurons. By contrast, successful neurons were scarcely found in the narrower group in judgement by either protocol I or II (green and magenta in **Fig. 4a**, respectively). The fraction in the narrower group of successful neurons was smaller than that of other neurons (*p* = 5.6 × 10^−7^ in protocol I, *p* = 1.7 × 10^−8^ in protocol II, χ^2^ test for a spike duration <0.5 ms), suggesting that our judgement scarcely makes mistakes in adopting fast-spiking neurons as projection neurons.

**Fig. 4.**
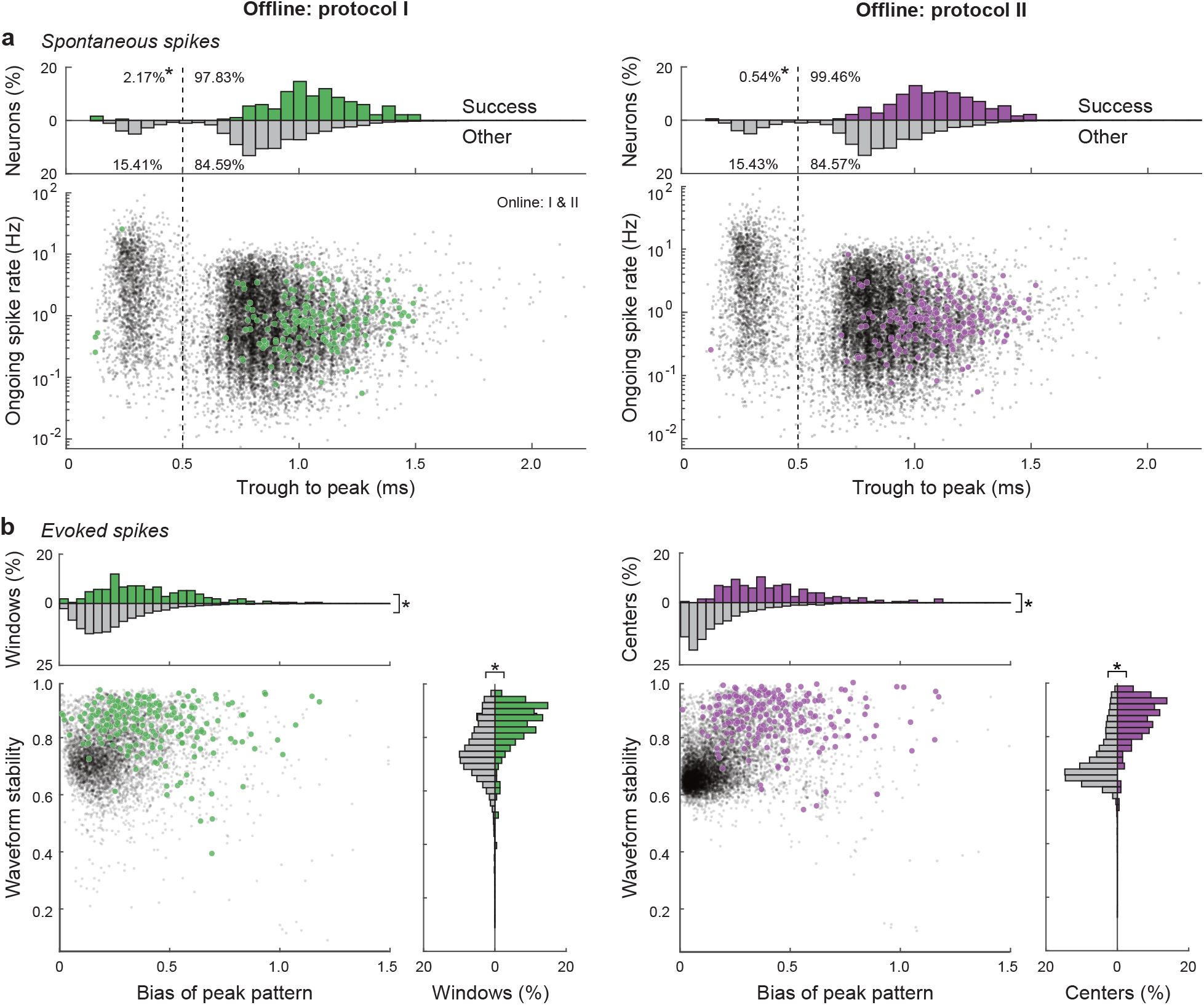
Comparison in spike properties between successes and others. **a**, Comparison in waveform durations of spontaneous spikes between successful (green and magenta dots) and other neurons (gray dots), in relation to subtypes of cortical neurons. We confirmed bimodal distributions (gray histograms in top) of the trough-to-peak durations of median spike waveforms and that the narrower type of cortical neurons exhibited higher ongoing spike rates than the wider type. Asterisks (*) mean *p* < 0.005 in 2×2 χ^2^ tests for the fractions with a trough-to-peak longer than 0.5 ms, success vs. other. **b**, Properties of inferred antidromic spikes to differentiate “success” from “other.” Each dot in the scatter plots represents the waveform stability and the peak pattern bias of an inferred group of antidromic spikes defined by the window (protocol I, left) and the center spike (protocol II, right) such that the jitter is sufficiently short (<0.25 ms). Asterisks (*) mean *p* < 0.005 in rank-sum tests.

We next examined the properties of inferred antidromic spikes to differentiate successes from others. Even though the jitter of inferred antidromic spikes was sufficiently short (<0.25 ms), the inference might not always be true. Actually, there were numerous sets of inferred antidromic spikes for which no neuron was found to pass the collision tests. We attempted to find properties to improve the inference of antidromic spikes. Because true antidromic spikes should originate in an identical neuron, spike waveform patterns on four channels of a tetrode should be stable in every stimulation trial. We evaluated the waveform stability on the basis of trial-by-trial deviation from the median waveform pattern. The waveform stabilities of the successfully identified antidromic spikes were higher than those of the others (vertical values in **Fig. 4b**; *p* = 7.1 × 10^−39^ in protocol I, *p* = 7.1 × 10^−70^ in protocol II, rank-sum test).

If synaptic spikes are evoked together in neuronal population around a tetrode, our inference of antidromic spikes by protocols I and II may adopt their population spike waves by mistake, which would exhibit an unbiased and stable peak pattern of four channels of the tetrode. By contrast, spike peak patterns of a single neuron would generally exhibit a bias depending on the direction of the neuron. Actually, we found that peak patterns of evoked spikes for successfully identified targets were more biased than those for the others (horizontal values in **Fig. 4b**; *p* = 4.9 × 10^−28^ in protocol I, *p* = 5.5 × 10^−67^ in protocol II, rank-sum test). Most of the unsuccessfully identified targets were distributed around the region of low bias and stability, whereas the successfully identified targets ones scarcely fell around this region. This property implies that the online *Multi-Linc* controller might be improved by excluding hopeless targets of collision tests.

We next examined the relationships between trigger and evoked spike waveforms (**Fig. 5**). We examined the similarity of the relative waveform patterns including several sampling points around the peak offline (vertical values in **Fig. 5a**). The relative waveform similarity between trigger and evoked spikes of the successful pairs was greater than that of the other pairs (*p* = 4.9 × 10^−114^ in protocol I, *p* = 4.6 × 10^−104^ in protocol II, rank-sum test). We confirmed the similarity of trigger and evoked spikes in the successful pairs on the basis of the details of the waveform patterns.

**Fig. 5.**
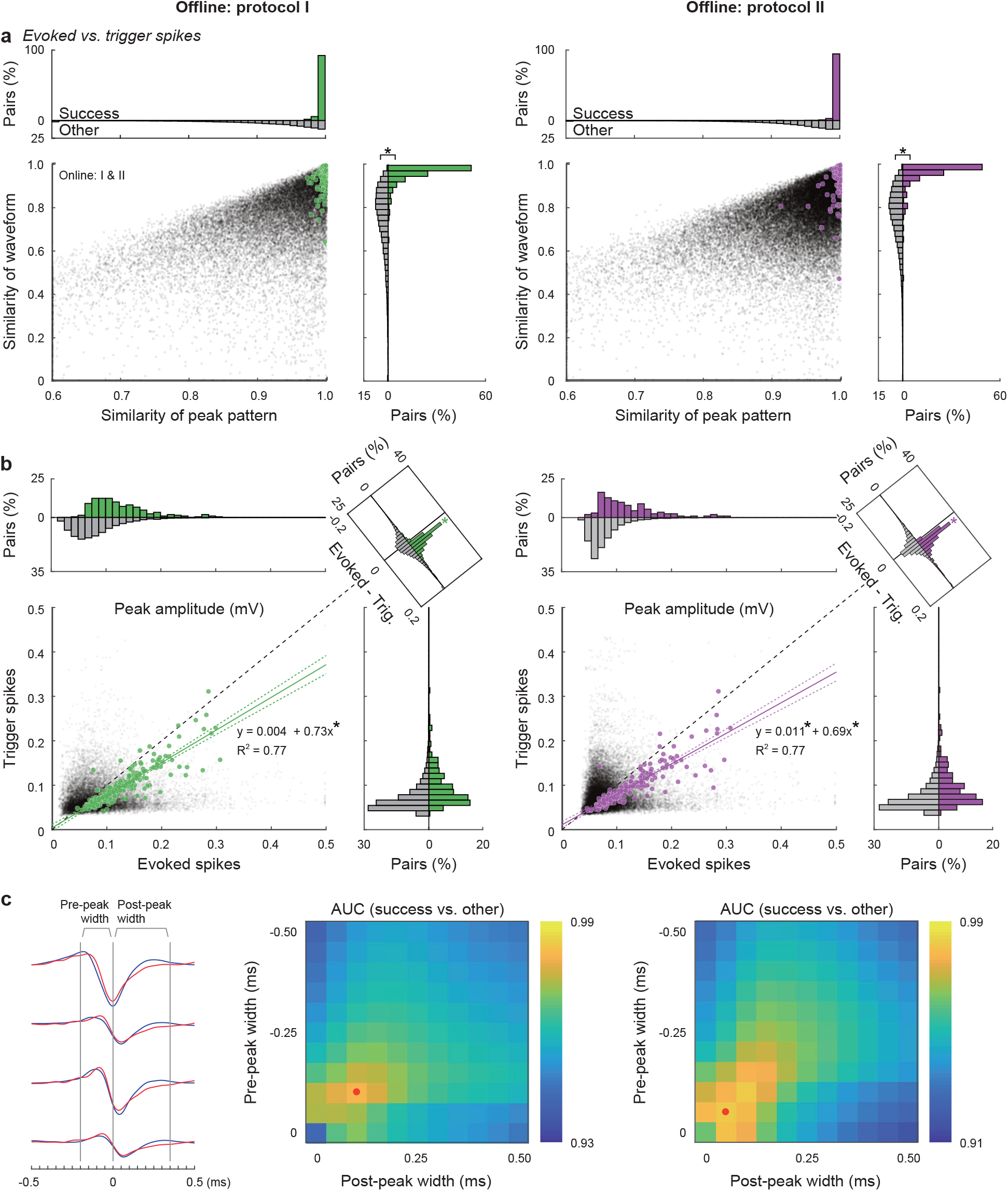
Similarity between trigger and evoked spike waveforms. **a**, Similarities in median waveforms between trigger and evoked spikes in the successful pairs in collision tests (green and magenta dots and histograms) and those in the other pairs (gray dots and histograms). Horizontal values represent the similarity for relative peak patterns on four tetrode channels (measured by direction cosines, common to the online controller), and vertical values represent the similarity for waveform patterns around peaks (−0.25 to +0.5 ms from the time point of the largest peak, 16 pt × 4 ch, measured by Pearson’s correlation coefficients; asterisks (*) mean *p* < 0.005 in rank-sum tests for success vs. other). **b**, Similarity between peak amplitudes of trigger and evoked spikes. The peak amplitude (largest of four channels) of trigger spikes in each successful pair was linearly correlated with that of evoked spikes (solid lines, linear regression; dashed curves, confidence intervals of 95%; asterisks (*) mean *p* < 0.005 in linear regression tests and signed-rank tests for trigger vs. evoked). c, Effective width of sampling points for waveform similarity to differentiate “success” from “other.” The combinations of pre-peak and post-peak widths (left) to calculate waveform similarity were swept in the range 0–0.5 ms (peak ± 0–10 sampling points). The AUCs in ROC analyses for the waveform similarity of the successful pairs against the others are shown as a function of the pre-peak and post-peak widths. The maximum AUCs (red dots) were obtained at the widths of a few points (middle, peak ± 2 pts in protocol I; right, peak ± 1 pt in protocol II).

Horizontal values represent the similarity of the relative peak patterns, which is the common index used to trigger collision tests in the online controller. The majority of successful pairs exhibited high peak-pattern similarity, confirming the condition used in the online controller (>0.99; 92% in protocol I, 94% in protocol II). This result suggests that the use of relative peak patterns worked well to trigger successful collision tests. However, successful pairs with lower peak-pattern similarity still existed. These pairs were picked by the offline analysis; they might not be targeted by the online controller. They also exhibited large similarity in waveform patterns; hence,the use of waveform patterns might improve the efficiency of the online controller.

Note that a substantial number of target pairs failed to pass collision tests even though the waveforms of trigger spikes and evoked spikes were similar (gray dots around [1,1] in **Fig. 5a**). These target pairs would be regarded as light-responsive neurons in the widely used “phototagging” technique. Although unsuccessful pairs might contain insufficient trigger trials, we could find examples of sufficient trigger spikes that failed collision (**Supplementary Fig. 6a**), providing a warning that a short jitter does not always correspond to direct antidromic activation of ChR2-positive neurons.

We next confirmed the relationships of the absolute amplitudes of trigger and evoked spikes. The amplitude of the evoked spikes on the peak channel was linearly correlated with that of trigger spikes in successful pairs (**Fig. 5b**; *p* = 0.29, *p* = 1.6 × 10^−6^8 for the zeroth and first orders in protocol I, *p* = 2.2 × 10^−3^, *p* = 3.7 × 10^−65^ for the zeroth and first orders in protocol II, *R*^*2*^ = 0.77 in both protocols, linear regression test), whereas the evoked spike sizes were larger than the trigger spike sizes (difference histograms in insets of **Fig. 5b**; *p* = 8.1 × 10^−33^ in protocol I, *p* = 1.2 × 10^−31^ in protocol II, signed-rank test). This result might be attributable to the fact that spontaneous spikes are just initiated in the depolarized soma with high synaptic conductance, whereas antidromic ones reach full size after axonal propagation into hyperpolarized soma. Irrespective of the cause, we can utilize this property to improve the online *Multi-Linc* controller. The amplitude information can be used to trigger the collision tests by shrinking spike waveforms of evoked spikes with a certain ratio.

The online *Multi-Linc* controller utilized only the spike peak patterns to trigger the collision tests. If spike waveforms around the peaks are also utilized, then the performance might be improved. The waveforms up to 0.5 ms after the peak can be incorporated in real-time processing. We evaluated the segregation between successful pairs and others in the correlation coefficient of spike waveforms incorporating various patterns of pre-peak and post-peak widths (**Fig. 5c**). The segregation was quantified by AUC of ROC analyses of the correlation coefficients. In judgement by either protocol I or II, the optimal segregation was obtained in waveform patterns within peak ± 0.15 ms and the waveform of peak ± 0.05 ms (three points of peak ± 1 at 20 kHz sampling) exhibited sufficient segregation (AUC = 0.98 in protocol I, AUC = 0.98 in protocol II). The use of waveform similarity of a few time points around the peak can improve the efficacy in triggering the collision tests.

One of the advantages in our *Multi-Linc* method is the ability to simultaneously identify multiple patterns of projections in a single recording session. To identify multiple projections in a session, we attempted to apply both recording and stimulation for bilateral motor cortices. We successfully obtained examples of reciprocal projections and projections to different target areas (**Fig. 6**). We also confirmed that the online *Multi-Linc* controller worked with virus injection to express ChR2 in neurons of a wild-type rat (**Supplementary Fig. 6b, c, d**).

**Fig. 6.**
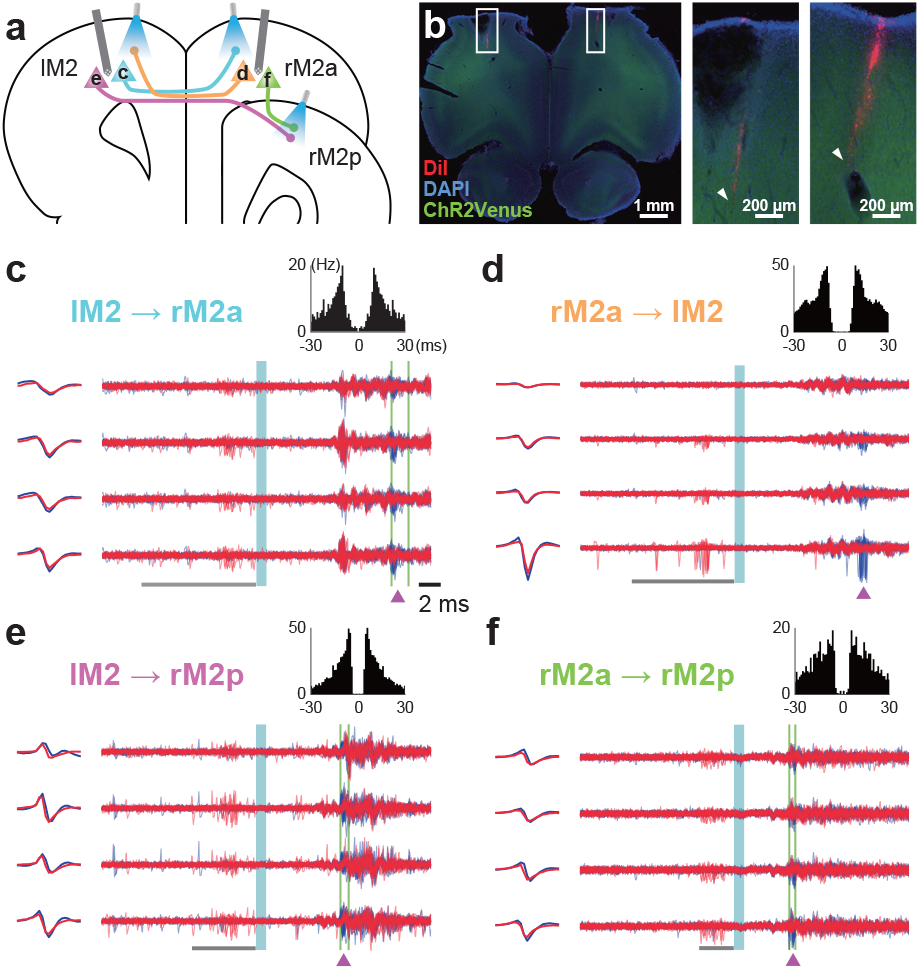
Simultaneous spike collision tests in reciprocally connected areas. **a**, Coronal view of bilateral recording and stimulation sites in the secondary motor cortex (M2). Neurons c–f correspond to panels **c–f** below, respectively. **b**, Histological confirmation of silicon probe insertion. The right two photos (magnified from the white boxes at left) indicate insertion tracks (arrowheads) marked with red fluorescent dye DiI (blue, DAPI; green, ChR2-Venus). **c–f**, Successful spike collision tests by the *Multi-Linc* system, which simultaneously identified four neurons projecting from left M2 (lM2) to right M2, anterior (rM2a) (**c**), from rM2a to lM2 (**d**), from lM2 to right M2, posterior (rM2p) (**e**), and from rM2a to rM2p (**f**). The corresponding data are shown in **Fig. 3a**.

## Discussion

Thus far, several methodologies have been used to examine functional spike signals between brain areas in behaving animals. Juxtacellular recording, a conventional electrophysiological technique (Pinault, 1996), enables spike measurement and post hoc visualization of a single neuron, which can be used to track its axonal projection histologically. However, juxtacellular neuron identification consumes large numbers of valuable task-trained animals because only a few neurons can be visualized histologically from each animal (Isomura et al., 2009; 2013).

Specific gene expression via retrograde viral vectors (Kato et al., 2011; Tervo et al., 2016) is useful in examining the functional activity of numerous projection neurons, especially when used in combination with two-photon laser-scanning or microendoscopic calcium imaging techniques (e.g., Osakada et al., 2011; Ren et al., 2018) with genetically encoded calcium indicators (GECIs) (Nakai et al., 2001; Zhao et al., 2011; Chen et al., 2013). Calcium imaging captures relative changes in the spike rate with low temporal resolution but not precise spike events. Hence, analyzing spike correlation with other neurons or with neural oscillations such as theta and gamma waves is almost impossible (Isomura et al., 2006; Sirota et al., 2008). Moreover, only a few projection pathways can be distinguished even by the latest multi-color calcium imaging techniques with different GECIs (Inoue et al., 2019) from retrograde viral vectors. In addition, the expression of GECIs takes at least several weeks after viral injection to the target areas.

In recent years, the “optotagging” (phototagging) technique has been widely used to identify neuron subtype or axonal projection by specifically stimulating ChR2-expressing neurons optogenetically (Jennings & Stuber, 2014; Silva et al., 2018; Starkweather et al., 2018). However, in the case where excitatory glutamatergic neurons are designed to express ChR2 for optotagging, unrelated ChR2-negative neurons could also respond to light stimuli synaptically through excited neurons (see **Fig. 1**). Such false-positive neurons cannot be excluded by optotagging alone even if spike jitter is sufficiently small (see **Fig. 3c and Supplementary Fig. 6a**). Thus, projection-specific expression does not always mean projection-specific excitation, which is why the spike collision test is still necessary for electrophysiological identification of axonal projection.

The greatest drawback of the spike collision test is its inefficiency. Here, we have paved the way to overcome this drawback by automating and parallelizing multi-channel spike collision tests. Our new *Multi-Linc* analysis has practical advantages over other methods in the exploration of projection neurons. Using transgenic animals expressing ChR2 broadly in the brain (Tomita et al., 2009; Saiki et al., 2018), we can flexibly test spike collisions in a different combination of multiple stimulating and recording sites in each experiment. The method is not restricted by the number of excitation wavelengths or by the sites and timing of viral injection. The method should also be easily applicable to long-distance pathways in the primate brain, which ensures sufficient spatiotemporal separation between stimulation and recording. Because of the versatility of the *Multi-Linc* analysis method, we expect to simultaneously observe cooperative or feedforward-feedback signaling through reciprocal projections between areas (see **Fig. 6**).

Nevertheless, our *Multi-Linc* analysis still has substantial room for improvement in its efficiency and accuracy. First, because ChR2 is expressed throughout the neuron, antidromic spike signals would be contaminated with noisy spikes by stimulating somata and dendrites of other neurons or a false projection could be identified because of incorrect stimulation of the midway of its axon. To avoid these problems, we developed a ChR2 variant that preferentially localizes at axonal terminals and that is available for spike collision tests (Hamada et al., 2021). Such ChR2 optimization for *Multi-Linc* analysis will lead to improvements in its temporal efficiency and spatial accuracy. Second, the inducibility of antidromic spikes depends on the properties of axons and the degree of ChR2 expression, which could result in selection bias of neurons for testing (Swadlow, 1998). Such bias will be eliminated by sufficient ChR2 expression in every axon and by local laser stimulation. Third, additional neurons would be assayed accurately by developing a real-time closed-loop system with high-density multi-channel probes (e.g., Neuropixels; Jun et al., 2017; Steinmetz et al., 2019). If spikes can be optically imaged in vivo using genetically encoded voltage indicators (Gong et al., 2015), an “electrodeless” *Multi-Linc* analysis could be developed in the future. Lastly, the algorithm for *Multi-Linc* analysis needs to be further improved to enhance its performance (e.g., to achieve precise detection of evoked spikes on the basis of the all-or-none law, optimal prioritization of neurons to be tested, and a stimulation protocol that does not interfere with brain states and behavior). These improvements could eventually lead to a “high-throughput” *Multi-Linc* analysis to elucidate the principle of spike communication among brain areas.

## Methods

### System for online *Multi-Linc* procedure

#### Hardware

We performed *Multi-Linc* experiments with a closed-loop controller for multi-channel recording and optogenetic stimulation. The online controller system was composed of a real-time processing computer [Intel Xeon Silver 4110 CPU @ 2.10GHz, 8 cores; Linux OS, Ubuntu 14.04, real-time kernel; D-TACQ Solutions, Scotland, UK] connected to a set box for input and output interfaces (D-TACQ ACQ2106; 128 channels of 16-bit analog inputs and 32 channels of digital-TTL outputs). In the system, data acquired through the 128 input channels at 20 kHz synchronous sampling were assured to transfer every 1.6 ms to the shared memory available on the CPU of the computer and arbitral TTL patterns of 20 kHz clocks set on the shared memory were assured to transfer every 1.6 ms to the output terminals. The analog input channels are supposed to receive amplified multi-channel signals from silicon probes inserted into the brain. The digital output channels are supposed to send signals to a multi-fiber laser stimulation device for optogenetic stimulation.

For analog inputs from multi-channel recording, we used 32-channel silicon probes (ISO_3x_tet_ A32; with seven tetrode-like electrodes on three shanks, NeuroNexus, Ann Arbor, MI). The signals from the silicon probes were amplified 2000 times and bandpass-filtered between 0.5 Hz and 10 kHz through pre- and main analogue amplifiers (MPA32I and FA32I, Multi Channel Systems, Reutlingen, Germany).

For optogenetic stimulation through digital TTL outputs, we used a multi-fiber laser stimulation device (MiLSS, custom-made; ASKA, Hyogo, Japan) to emit a blue light pulse (445 nm, 7–15 mW at the end) into each of seven optical fiber ports with a two-axis galvanometer mirror under the control of 8-channel TTL signals. The light pulse was delivered after the laser stimulation device started emitting the laser beam (1 ms delay) and the galvanometer mirror moved (as long as 0.75 ms).

The multi-channel inputs and TTL outputs were bifurcated to sets of 32-channel digital recording devices (LX-120, TEAC, Tokyo, Japan; 16 bit, 20 kHz) for the offline *Multi-Linc* procedure after the experiments.

### Software

We composed the controller software of a web-based user interface (UI) by CGI (Common Gateway Interface; Perl 5, lighttpd 1.4) and process commands implemented in the C and C++ languages controlled by shell scripts. Experimenters can manipulate the system through standard web browsers on the local network by setting parameters and sending start and stop signals for the respective procedures. The commands for start and stop signals are called by CGI form. Once the main process starts, multiple commands run and interact in real time through the shared memory.

### Real-time filtering and spike detection

We implemented simple high-pass filtering for real-time spike signal detection with a one-side exponential filter:

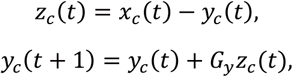

where x_c_ (t) is the local field potential acquired from channel c at time point t, y_c_ (t) is the local average, and z_c_ (t) is the filtered value. The parameter G_y_ determines the averaging scale (default 0.25 ms corresponds to G_y_ = 0.2 at 20 kHz sampling). Spike detection was based on the noise level on a tetrode of four channels. To avoid square-root computations at every step, we compared the square summation of four channels with the noise variance. Because a spike signal appears to be a negative sharp peak in extracellular recoding, we considered only the negative components,

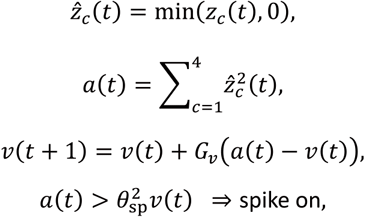

where a(t) is the square amplitude of the rectified four-channel vector of a tetrode and v(t) is an estimate of the amplitude variance. The parameter G_v_ determines the averaging scale of the variance estimation (default 1 sec corresponds to G_v_ = 5 × 10^−5^). The parameter θ_sp_ is the threshold of spike detection relative to the noise level (θ_sp_ = 4 [SD] at default). The peak time point in the range of successive “spike on” is defined as the spike time t_k_, and the four-channel vector at the time, 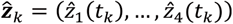 and the squared amplitude a_k_ = a(t_k_), are stored in the shared memory. The real-time filtering and spike detection always run for every tetrode parallelly from the session start to the end.

#### Evoked response collection

A stimulation site is serially selected and the stimulation signal is sent through the shared memory at a random timing such that inter-stimulation intervals might be longer than the criterion (i.e., the minimum interval between the same-site stimulations, Isame) and between the different-site stimulations, Idiff (Isame = 1.0 sec, Idiff = 0.5 sec at default). These minimum intervals are set to avoid epileptic neural responses. The duration of a stimulation (TTL on) is set via parameter Di for each stimulation site i (default: 1.0 ms). Experimenters can manipulate the duration of each site through pre-session tests. After every stimulation, raw data {x_c_ (t)} from all tetrodes during a time range Trange (default: 30 ms) before and after the stimulation are stored from the ring buffer on the shared memory. While waiting the minimum inter-stimulation intervals, the stored {x_c_ (t)} data are filtered semi-offline with a precise symmetric filter common to the offline analyses described in the section of the offline *Multi-Linc* procedure. The high-pass filter is designed to subtract Gaussian-smoothed signals (σ = 0.25 ms).

The precisely filtered data are stored in the files for use in the next process. The process of the evoked response collection stops so the next process can be started when the required number of stimulations per stimulation site, Nanti (default: 100) have been collected.

#### Antidromic spike inference

After the evoked responses are collected, the process of antidromic spike inference starts semi-offline. The precisely filtered evoked responses of each tetrode are normalized into the Z-score based on the estimated noise level during the range Trange before stimulations. The noise level is estimated for each tetrode commonly among four tetrode channels by averaging over the stimulation trials of all the stimulation sites. Antidromic spikes in evoked responses are inferred for each tetrode–stimulation site pair. We used two different inference protocols (protocols I and II, see **Fig. 2**). The inference protocols were common to the offline analyses. The details of the inference protocols are described in the section of the offline *Multi-Linc* procedure. The averaged peak amplitude patterns of inferred antidromic spikes are transferred to the next process.

During the semi-offline process for the antidromic spike inference, frequency-following tests can be optionally executed as a parallel process. Several repetitive stimulations with short intervals are executed for every stimulation site. Because ChR2 itself cannot respond at such a short interval (Gunaydin et al., 2010), the frequency-following test is not effective to verify the collision test in the present experimental paradigm. Therefore, we implemented it as an option and confirmed that it could work (**Supplementary Fig. 6e**).

#### Collision test

The process of the collision test monitors spike occurrence on all tetrodes in real time through the shared memory. When a spike is found on a certain tetrode, the targets associated with the tetrode and the stimulation sites such that the intervals from the last stimulations are longer than the criteria of the minimum inter-stimulation intervals, Isame and Idiff, are sought. The squared direction cosine to each targeted peak pattern is calculated as the similarity to the inferred antidromic spikes, 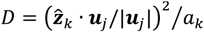, where u_j_ is the peak pattern vector of the j-th target for collision tests. If the similarity is greater than the criterion, D>θ_trig_^2^ (default: θ_trig_ = 0.99), then the spike triggers a collision test for the j-th target and the stimulation signal of the corresponding site is immediately set in the shared memory. Packet communication every 1.6 ms with the input–output interfaces assures a TTL onset within 3.2 ms after the spike. If the number of collision tests for a target reaches the criterion, Ntest (default: 200), then the target is excluded. The process stops if all targets are excluded, or if the experimenter sends the stop signal.

### Experimental data collection

#### Animal preparation

All experiments were approved by the Animal Research Ethics Committee of Tamagawa University (Animal Experiment Protocol H30-32) and the Institutional Animal Care and Use Committee of Tokyo Medical and Dental University (A2019-274) and were carried out in accordance with the Fundamental Guidelines for Proper Conduct of Animal Experiment and Related Activities in Academic Research Institutions (Ministry of Education, Culture, Sports, Science and Technology of Japan). All surgeries were performed under appropriate isoflurane anesthesia, and all efforts were made to minimize animal suffering. The procedure for animal experiments was established in a series of studies by our group (Isomura et al., 2009, 2013; Soma et al., 2017; Nonomura et al., 2018; Saiki et al., 2018; Rios et al., 2019; Kawabata et al., 2020). We used eight adult male rats (407.6 ± 32.2 g; six Wister Thy1.2-ChR2V4 line backcrossed with the Long–Evans strain (Tomita et al., 2009; Saiki et al., 2018); two wild-type Long–Evans strain (Sankyo Labo Service, Tokyo, Japan) for viral vector injection). These rats were kept under an inverted light schedule (lights off at 12 AM; lights on at 12 PM) in their home cages to adapt to experimental surroundings.

#### Surgery

Primary surgery was performed to attach a head-plate (CFR-2, Narishige, Tokyo, Japan) to the skull of rats under anesthesia by isoflurane gas (4.0– 4.5% for induction and 2.0–2.5% for maintenance, Pfizer Japan, Tokyo, Japan) using an inhalation anesthesia apparatus (Univentor 400 anesthesia unit, Univentor, Zejtun, Malta). The body temperature was maintained at 37.0°C using an animal warmer (BWT-100, Bio Research Center, Aichi, Japan) during anesthesia. The head of rats was fixed on a stereotaxic frame (SR-10R-HT, Narishige) with ear bars, and applied with lidocaine (Xylocaine Jelly, Aspen Japan, Tokyo, Japan) for local skin anesthesia and povidoneiodine disinfectant solution (10%, Kaneichi, Osaka, Japan) for disinfection around surgical incisions. The head-plate was then glued to the skull with stainless steel screws and dental resin cement (Super-Bond C & B, Sun Medical, Shiga, Japan; Unifast II, GC Corp., Tokyo, Japan), and reference and ground electrodes (PFA-coated silver wires, A-M systems, WA; 125-mm diameter) were implemented under the bone on the cerebellum. Analgesics and antibiotics (meloxicam, 1 mg/kg sc, Boehringer Ingelheim, Tokyo, Japan; gentamicin ointment, 0.1% us. ext., MSD, Tokyo, Japan) were finally applied to remove pain and prevent infection.

More than 1 week later, secondary surgery was performed to make cranial windows to frontal motor cortices bilaterally (1.0–3.5 mm anterior and ±1.0–3.0 mm lateral from the bregma) under the isoflurane anesthesia. The bone and dura mater were opened and removed by a dental drill (Tas-35LX, Shofu, Kyoto, Japan) and a dura picker (DP-T560-80, Bio Research Center, Aichi, Japan). The cortical surfaces were washed with PBS containing antibiotic (0.2% amikacin sulfate, Sawai, Osaka, Japan) and covered with antibiotic ointment (Chlomy-P ointment AS, Daiichi Sankyo Healthcare, Tokyo, Japan) and dental silicone sealant (DentSilicone-V, Shofu, Kyoto, Japan) until recording experiments.

#### Multi-channel recording and optogenetic stimulation

We performed online *Multi-Linc* experiments in the frontal motor cortices of unanesthetized rats under head-fixation. For the multi-channel recording, we used two 32-channel silicon probes, although the system can accommodate 128 channels. Approximately 1 h before each recording session, the probes were inserted to a depth of 1.0–1.5 mm from the cortical surface, typically in layer 5, where intratelencephalic (IT)-type projection neurons are distributed most abundantly, using three-axis micromanipulators (SMM-200B and SMM-100, Narishige). On the last recording day, the probe tracks were marked with the red fluorescent dye DiI (DiIC18(3), PromoKine, Heidelberg, Germany) applied to the back of each shank for histological conformation. In the optogenetic stimulation, we used micromanipulators (SM25A, Narishige) to place two to six optic fibers (FT1000EMT, diameter: 1000 μm, Thorlabs, New Jersey, USA) on the cortical surface in a symmetrical position contralaterally from the silicon probes.

#### Histology

After all recording sessions, the rats were perfused transcardially with cold saline and subsequent 4% formaldehyde in 0.1 M phosphate buffer under deep anesthesia with urethane (3 g/ kg, ip, Nacalai Tesque, Kyoto, Japan) to confirm the final probe tracks. The brains were removed and post-fixed at least overnight, and 50 μm-thick serial coronal sections were prepared using a microslicer (VT1000S, Leica, Wetzlar, Germany). The serial sections were cover-slipped with mounting medium (DAPI Fluoromount-G, Southern Biotech, AL, USA) and observed under a fluorescence microscope (IX83 inverted microscope, Olympus, Tokyo, Japan).

### Offline *Multi-Linc* procedure

#### High-pass filtering and spike sorting

Signals from each tetrode channel were filtered with a high-pass filter designed to subtract Gaussian smoothed signals (σ = 0.25 ms) throughout an experimental session (several hours). The filtered data were normalized into Z-scores on the basis of the common noise level estimated for each tetrode among four tetrode channels throughout the session. Spike events of individual neurons were isolated and clustered in each tetrode using the automatic spike sorting software EToS (Takekawa et al., 2012) and the manual spike clustering software Klusters (Hazan et al., 2006; **Supplementary Fig. 2d**).

#### Antidromic spike inference in protocol I

Filtered responses on a certain tetrode evoked by stimulations to a certain site were aligned with the onset of stimulation. The temporal window of a fixed width (2 ms width in the online experiments, tuned to 1 ms in the offline analyses) was slid along time from channel 1 to 4 (**Fig. 2b**, top), and the negative peak within the window was detected in each trial (**Supplementary Fig. 2a**). For each window, the distribution of the negative peaks was obtained (dots in **Fig. 2b**, middle). The goodness of the window can be characterized by the first quartile (75% from the largest) of the amplitude distribution because a larger amplitude quartile indicates a more stable occurrence of spikes within the window (**Supplementary Fig. 2b**). All the windows of timings and channels were characterized in the same manner. The windows with the amplitude quartiles below a certain threshold were excluded from candidates. Among the remaining candidates, the window with the maximum amplitude quartile was adopted as the best window (**Fig. 2b**, bottom, and **Supplementary Fig. 2c**). Once adopted, the window width was fitted to the jitter distribution. Using 75% from the largest peak, the jitter was estimated as the quartile deviation QD of the peak timings. The window start and end were reset by multiplying a factor Wjitter to the deviation from the median timing as median ± Wjitter QD (Wjitter = 5, online; Wjitter = 4, offline). The distribution of negative peaks within the new window was determined again, and the window start and end were reset in the same manner. If the window converged after the iteration, we adopted the window to determine inferred antidromic spikes. We determined 75% from the largest peak within the converged window as representative spikes inferred to be antidromic. The median waveform of each tetrode channel was calculated among the representative spikes at each time point relative to the peak timings. After an inferred window was adopted, the sliding windows on all channels overlapping the time range of the adopted window were excluded from candidates, and the selection of an inferred window among the remaining candidates was iterated.

#### Antidromic spike inference in protocol II

The protocol II evaluated the aggregation of timings and waveform features of spikes. First, spikes in evoked responses were detected by thresholding (negative peaks below 5 SD). We defined the similarity between two evoked spikes as

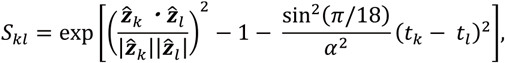

where z^_k_ and t_k_ are the four-dimensional vector of the k-th spike amplitude pattern on the tetrode channels and the timing from the stimulation onset, respectively. Spike amplitudes were rectified to be non-positive values, and the angle difference of two vectors was smaller than π/2; hence, the similarity decreased monotonically with increasing angle difference. The similarity of identical spikes gives the maximum value S_kk_ = 1. The parameter α controls the weight of the timing deviation. The timing deviation t_k_ - t_l_ = α corresponds to the angle difference of π/18 between peak amplitude directions. The acceptable jitter α was tried in the range 0.1–4 ms for the online controller and 1 ms for the offline analyses.

We searched the center of aggregated spikes among all trials of stimulations on the basis of the aforementioned similarity. We focused on each detected spike as a candidate for the aggregation center and determined the spikes most similar to the focused spike in respective trials (**Supplementary Fig. 2a**). The degree of aggregation around the focused spike can be characterized by the first quartile in the similarity distribution (75% from the most similar one; **Fig. 2c and Supplementary Fig. 2b**). All detected spikes were characterized by the similarity quartiles in the same manner. Spikes with similarity quartiles less than a threshold θ_aggr_ were excluded from candidates for the center spike (online default: θ_aggr_ = 0.98; candidates were not restricted in the offline analyses to gather control samples). Among the remaining candidates, the center spike with the maximum similarity quartile was adopted as the aggregation center (**Fig. 2c, bottom, and Supplementary Fig. 2c**). As in protocol I, we used 75% of the nearest spikes to the adopted center spike as representative spikes inferred to be antidromic. The median spike waveform and the median timing were calculated among the representative spikes. Once adopted, 75% of the nearest spikes were excluded from candidates for the next center spike. Selection of the center spike was then iterated.

#### Judgement in collision tests

In offline analyses, we tested all the pairs of neurons (spike clusters) and inferred sets of antidromic spikes (defined by windows in protocol I, and center spikes in protocol II) for whether spike collisions might occur. Two criteria were used for the determination (**Fig. 3c and Supplementary Fig. 1a**). The first criterion was based on the receiver operating characteristic (ROC) analyses to evaluate the elimination of evoked spikes by collisions. The second criterion was the jitter of inferred antidromic spikes.

For each pair of a neuron and an inferred set of antidromic spikes, the trigger and no-trigger stimulation trials were defined. When spikes of the neuron occurred within 2 ms before the minimum latency of the inferred antidromic spikes (the representative spikes defined in either protocol I or II), the stimulation trial was excluded for analyses because it could not determine whether the evoked spike elimination might originate in spike collision or neuronal refractory. From the remainder of the stimulation trials, we extracted trigger trials in which spikes of the neuron occurred in the trigger range before the stimulations. The trigger range was defined such that a spontaneous spike within the range should collide with the antidromically evoked spike (see **Supplementary Fig. 1b**). If the number of the extracted trigger trials was less than 15, then the focused pair was excluded from the targets of collision tests.

If the trigger trials were defined, then we extracted no-trigger trials in which no spikes of the neuron occurred in the no-trigger range before the stimulations from the trials near the trigger trials. The no-trigger range was defined such that a spontaneous spike within the range could collide with the antidromically evoked spike (see **Supplementary Fig. 1b**). We extracted the nearest 10 no-trigger trials per each trigger trial, and merged trials duplicated among different trigger trials. Thus, the number of extracted no-trigger trials was ten times the number of trigger trials at maximum.

We evaluated the elimination of spikes to be evoked with negative peak amplitudes within the inferred window in protocol I and the similarities to the inferred center spike in protocol II (Supplementary Figs. 1c,d and 2e,f). The ROC analysis was performed between distributions of the evaluated variables (peak amplitudes or similarities) in the extracted trigger and no-trigger trials, and the area under the ROC curve (AUC) was calculated. The criterion for spike elimination was defined on the basis of the AUC distribution among the whole tested pairs such that AUC might be more than five times the standard deviation σ robustly estimated with the relation in the Gaussian distribution, σ = median[|AUC -median [AUC ]|]/0.6745. The jitter was defined as the quartile deviation of the latencies of evoked spikes determined by the inferred window (protocol I) or center spike (protocol II) in no-trigger trials. Notably, the latency of each spike was calculated by spline interpolation for its waveform.

If AUC > 5σ and jitter < 0.25 ms, then the tested pair was defined to be successful in the collision test. If multiple successful pairs were found for an identical neuron, then only the pair with the highest AUC was adopted as a successful pair.

#### Analysis for the spike property of identified projection neurons

##### Spontaneous spikes

We computed the trough- to-peak durations of unfiltered and interpolated waveforms and the ongoing spike rate of spontaneous spikes for each neuron in the range from the first to the last stimulation in a session, except for a 30 ms period following each stimulation (**Fig. 4a**). The neurons belonging to successful pairs were defined as successful neurons. The other neurons were found to be unsuccessful for either set of inferred antidromic spikes.

##### Evoked spikes

We computed waveform stability and bias of the peak pattern for each evoked spike (**Fig. 4b**). The waveform stability was evaluated on the basis of the Pearson’s correlation of waveforms between the median waveform and the waveform in each no-trigger trial. The waveforms were extracted from −0.25 ms to 0.5 ms around the peak amplitude of spikes. The waveform stability was defined as the median of the Pearson’s correlation coefficients among the stimulation trials. The bias of the peak pattern was calculated as the coefficient of variation among the median peak amplitudes of the four channels of evoked spikes in the no-trigger trials.

##### Relationship of evoked spikes and trigger spikes

We evaluated the similarity of peak patterns, the similarity of waveforms, and the peak amplitudes for each pair of evoked spikes and trigger spikes (**Fig. 5a**,**b**). The similarity of a peak pattern was calculated as the direction cosine used in the online *Multi-Linc* procedure. The similarity of the waveform was calculated as the Pearson’s correlation coefficient of the median waveforms from −0.25 ms to 0.5 ms around the peak amplitude of the spikes. The peak amplitude was extracted from the channel with the maximum peak amplitude of the median waveform among four channels. In addition, we computed the discriminability (i.e., the AUC) between “success” and “other” pairs on the basis of the waveform correlation between the evoked and triggered spikes while varying the width of the waveforms used (**Fig. 5c**). The range of the waveform was varied to 0.5 ms before and after the peak (i.e., the pre-peak range and the post-peak range, respectively).

### Statistics

The details of the statistical tests are summarized in **Supplementary Table 2**.

## Code availability

The source codes of the commands described in c, c++, perl and shell-script to control the online *Multi-Linc* procedure will be available in GitHub (https://github.com) before publication.

## Data availability

Links for publicly available datasets will be provided before publication.

## Acknowledgements

We are grateful to Drs. S. Hamada, R. Hira, M. Kimura, K. Kobayashi, A. Nambu, S. Nonomura, T. Ohtsuka, A. Ríos, A. Saiki-Ishikawa, T. Sakairi, S. Soma, A.M. Watabe, and T. Yoshizawa for their helpful comments. We also thank M. Goto, M. Ikushima, R. Mizuno, T. Shimada, M. Takahashi, C. Soai, and H. Yoshimatsu for technical assistance.

This work was funded by Brain/MINDS JP19dm0207089 (Y.I.) from AMED; CREST JPMJCR1751 (Y.I.) from JST; Grants-in-Aid for Scientific Research on Innovative Areas JP16H06276 (Y.I.), JP18H05524 (Y.S.), and JP20H05053 (Y.I.) from MEXT; Scientific Research (B) JP19H03342 (Y.I.) from JSPS; a Start-up JP21K20684 (M.K.) from JSPS; and a research grant from Takeda Science Foundation (Y.I.).

## Author contributions

Y.I. and Y.S. designed the whole study. K.M., M.K., and Y.S. developed the system. M.K. collected the experimental data. K.M., M.K., and Y.S. analyzed the data. K.M., M.K., Y.I., and Y.S. wrote, discussed, and edited the manuscript.

## Additional information

Extended data is available for this paper at https://doi.org/***.

Supplementary information is available for this paper at https://doi.org/***.

**Fig. S1.**
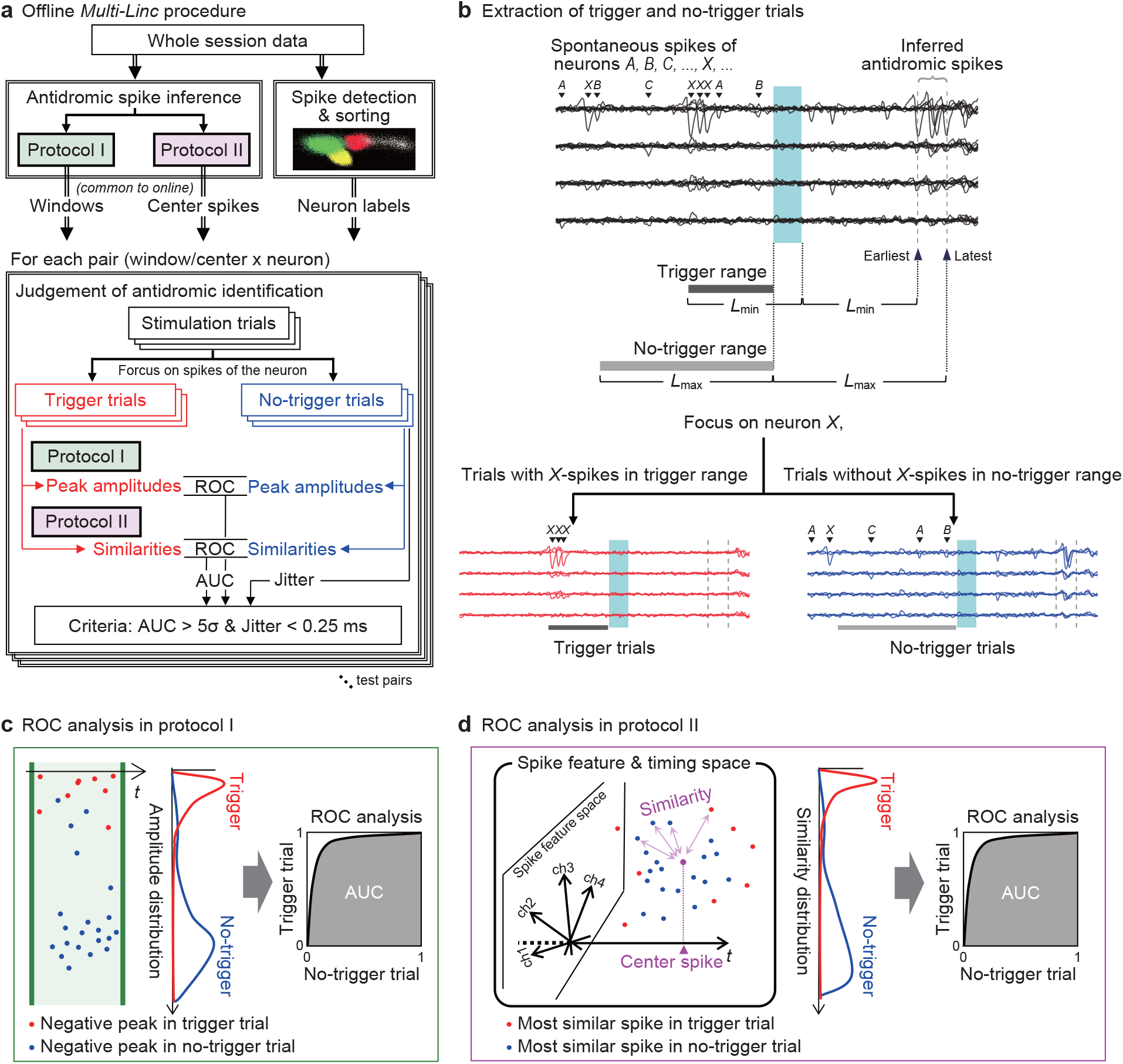
Procedure for the offline analysis of the collision test. **a**, Flow of offline *Multi-Linc* procedure. **b**, Procedure for extraction of trigger and no-trigger trials focusing on a certain pair of a neuron and a set of inferred antidromic spikes. We defined the trigger/no-trigger range (dark-gray/light-gray bar) before the stimulation such that spontaneous spikes in the range should/could collide with the evoked antidromic spikes. The possible minimum latency is inferred as the length from the offset of the stimulation light to the earliest timing of the inferred antidromic spikes, Lmin. Hence, a spontaneous spike Lmin before the light offset should not yet reach the axon terminal until the light offset and should collide with the evoked antidromic spike. We defined the trigger range from Lmin before the light offset to the light onset. However, the possible maximum latency is inferred as the length from the light onset to the latest timing of inferred antidromic spikes, Lmax. Hence, a spontaneous spike Lmax before the light onset might not yet reach the axon terminal at the light onset and might be in time to collide. We defined the no-trigger range of length Lmax before the light onset. If spontaneous spikes do not occur in the no-trigger range, then the collision should not occur. In these ranges before optical stimulations, multiple spontaneous spikes of multiple neurons (A,B,C, … X, …) occurred. Focusing on a neuron X, we extracted the stimulation trials with some spikes of X in the trigger range (dark-gray bar) as the trigger trials (red). Conversely, we extracted the trials without any spikes of X in the no-trigger range (light-gray bar) as the no-trigger trials (blue). Moreover, the no-trigger trials were restricted so that the terms of the trigger and no-trigger trials in the recording session might be equivalent for the sake of fair comparison (see **Online Methods**). **c, d**, Schema of ROC analyses in protocols I and II to judge whether the pair succeeds in the collision test. Elimination of antidromic spikes inferred to be evoked can be judged on the basis of the variables used in the antidromic spike inference: the negative peak amplitudes in protocol I (**c**) and the similarity to the center spike in protocol II (**d**).

**Fig. S2.**
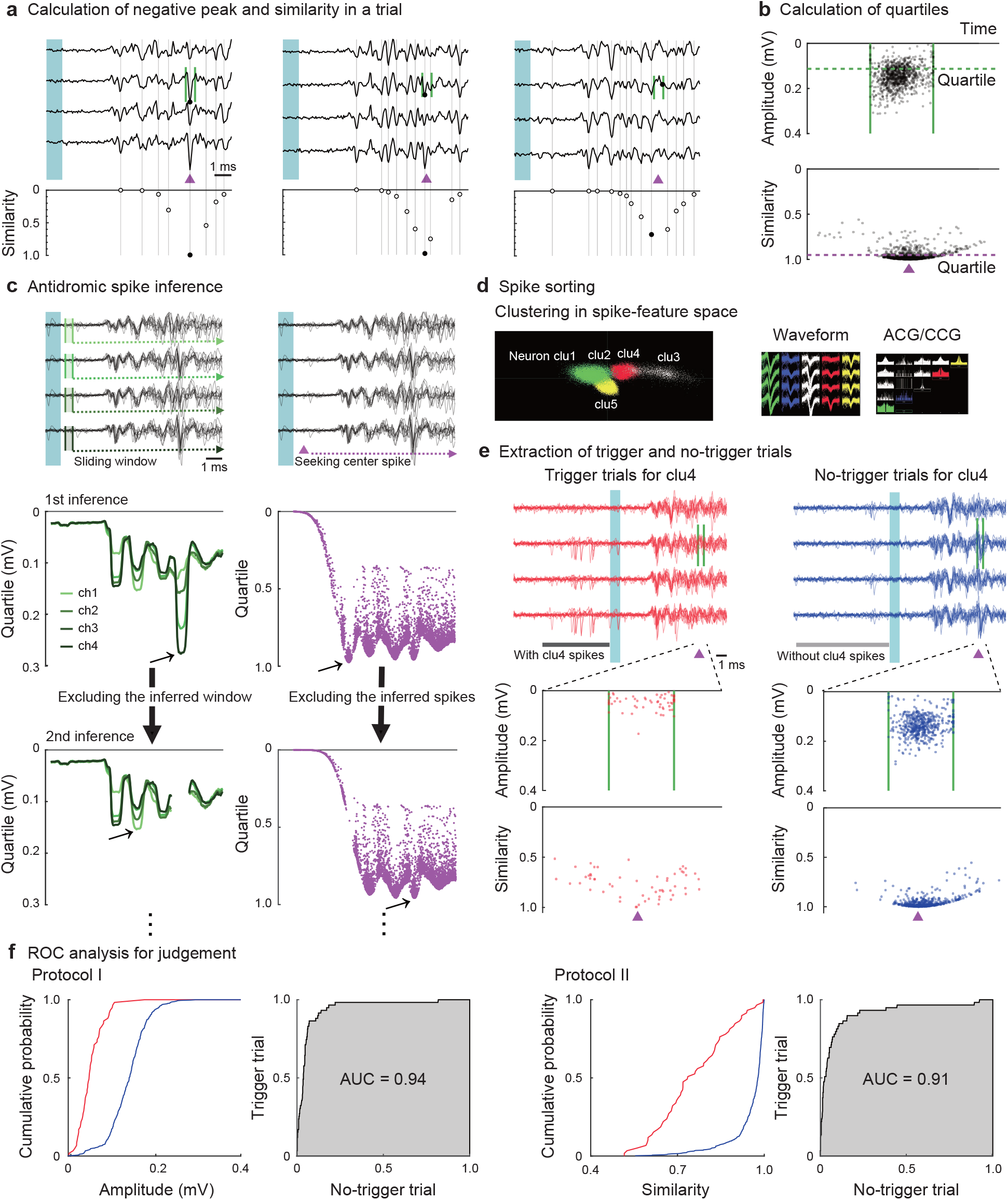
Examples of the offline analysis. **a**, Negative peaks (protocol I) and similarities (protocol II) in an example stimulation trial. The negative peaks in a window (green vertical lines) are shown as black dots in each trace. The timing of the focused center spike in a different trial is indicated by a magenta triangle. The similarities in timings and relative peak patterns of evoked spikes in this trial to the center spike are shown in the lower panels. The similarity of the most similar spike to the center spike is adopted in each trial (black filled dot). The highest similarity in a trial gives a high value if an evoked spike exists such that both the timing and relative peak pattern are similar to the center spike (left, center), and gives a low value otherwise (right). **b**, Distributions (scatter dots) and their lower quartiles (75% from the largest; dashed lines) of negative peak amplitudes within a window and highest similarities to a center spike in an example session. Each dot shows the negative peak amplitude (protocol I) and the highest similarity (protocol II) adopted in each trial, plotted against the spike timing. A high quartile value indicates that antidromic-like spikes are stably evoked in stimulation trials. **c**, Examples of antidromic spike inference. Antidromic spike inferences were performed on the basis of the quartiles of peak amplitudes (protocol I) and similarities (protocol II). The quartiles were calculated as functions of all sliding windows (green curves) or all seeking center spikes (magenta dots). The window or the center spike with the maximum quartile (black diagonal arrows) was adopted as the best inference of antidromic spikes. The spikes over the quartile were adopted as a set of inferred antidromic spikes. After the window range or the inferred spikes were excluded, the following window or center spike was searched. **d**, An example of spike sorting. All spontaneous spikes in a session were sorted into clusters likely to originate in identical neurons by a standard spike sorting technique (EToS; see **Online Methods**) to prepare for extraction of trigger and no-trigger trials. The plots show clusters in spike feature space (left), their overlayed waveforms (center), and ACG/CCG (right). **e**, Extraction of trigger and no-trigger trials for an example pair of a spike cluster (clu4, a putative identical neuron) and an inferred antidromic window (green vertical lines) or center (magenta triangles). For each cluster, trials with spontaneous spikes of the cluster in the trigger range (dark-gray bar; see **Supplementary Fig. 1b**) before stimulation were extracted as the trigger trials (top left, red). Trials without any spontaneous spikes of the cluster in the no-trigger range (light gray bar) were extracted from trials near the trigger trials as no-trigger trials (top right, blue). Negative peak amplitudes within the window (protocol I; middle) and similarities to the center spike (protocol II; bottom) were calculated for the respective sets of trigger (left) and no-trigger (right) trials. The lower value of the peak amplitude or the similarity in each trial indicates that an antidromic spike is more likely to be eliminated or not be evoked. If the peak amplitudes or the similarities in the trigger trials are sufficiently lower than those in the no-trigger trials, then spontaneous spikes of the cluster just before stimulations are likely to collide with the evoked antidromic spikes. **f**, Example of ROC analysis for judgement of whether the spike collision might occur. Cumulative distributions of peak amplitudes (protocol I; left half) or similarities (protocol II; right half) in trigger trials (red) and no-trigger trials (blue) and the ROC curves between trigger and no-trigger trials (black lines) are shown. The ROC curve represents the cumulative fraction at a certain value in the trigger trials as a function of that in the no-trigger trials. A larger area under the ROC curve (AUC; gray area) means a larger segregation between the trigger and no-trigger trials. The success of the collision was determined by a sufficiently large AUC (see **Fig. 3c**).

**Fig. S3.**
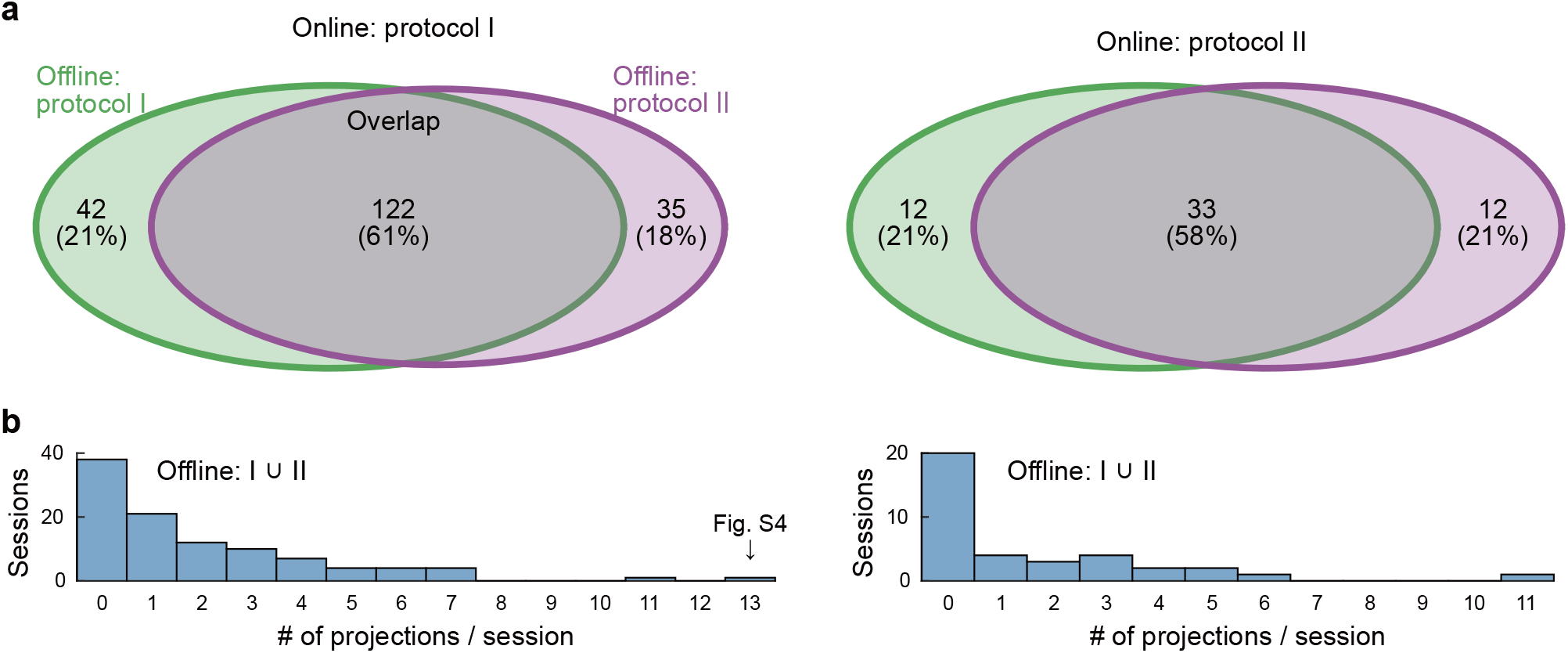
Overlap in identified neuronal projections by protocols I and II. **a**, Venn diagrams of the number of identified neuronal projections in the offline judgements by protocols I and II, separately shown for the experimental sessions obtained by using the online controllers with protocols I and II (left and right, respectively). The numbers and percentages (parentheses) are described in the respective regions. **b**, Histograms of the merged outcomes of identified neuronal projections per session by protocols I and II. The maximum merged outcome (13 projections) was obtained in a session using the online controller with protocol I, the details of which are shown in **Supplementary Fig. 4**.

**Fig. S4.**
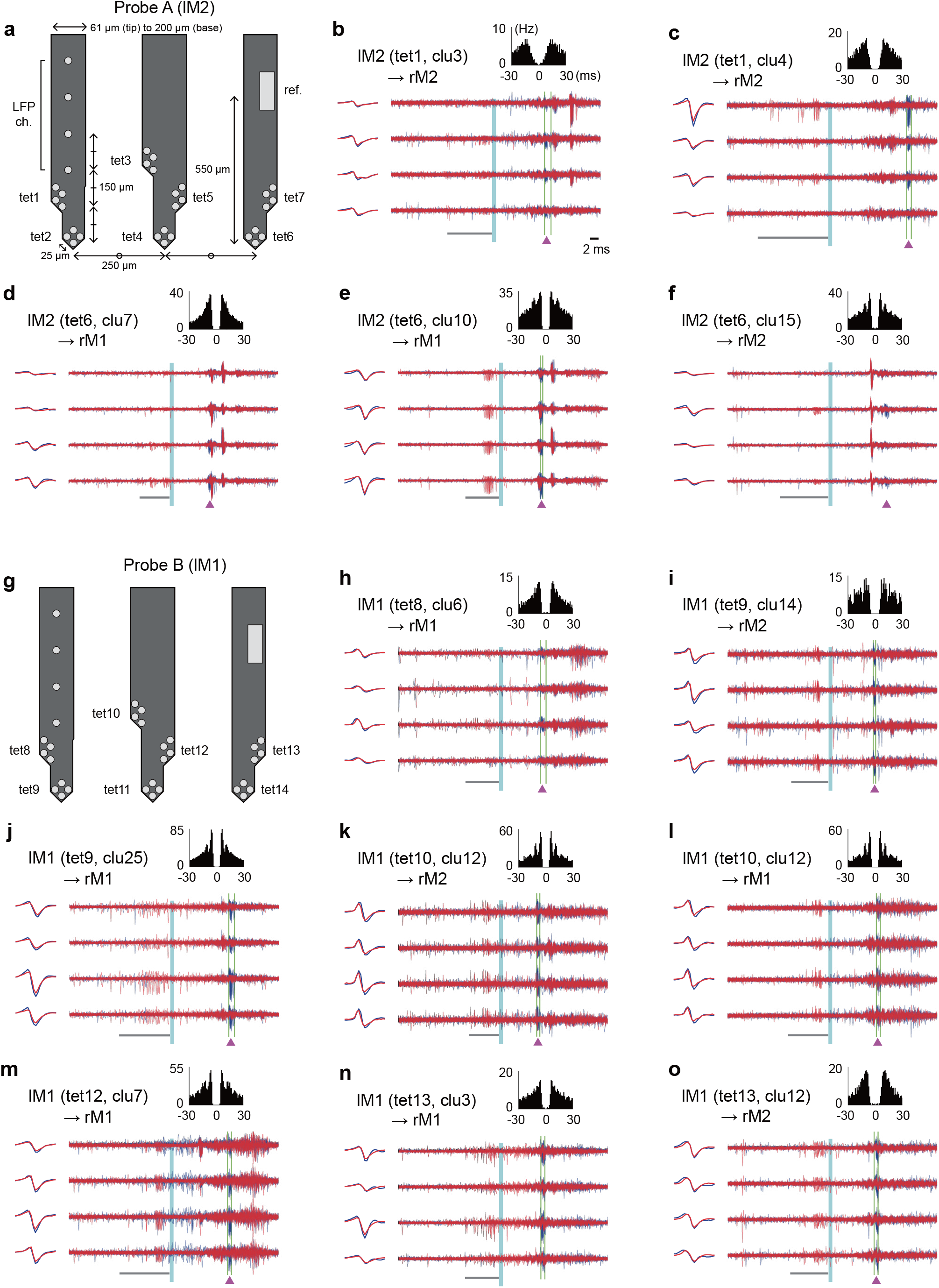
Thirteen successful spike collision tests in a session. **a** and **g**, Electrode arrangement of silicone probes A and B. **b–f**, Successful spike collision tests using the *Multi-Linc* system, which simultaneously identified five projection neurons recorded from left M2 (probe A). **h–o**, Successful spike collision tests that identified eight projection neurons in left M1 (probe B). Data are shown in **Fig. 3a**.

**Fig. S5.**
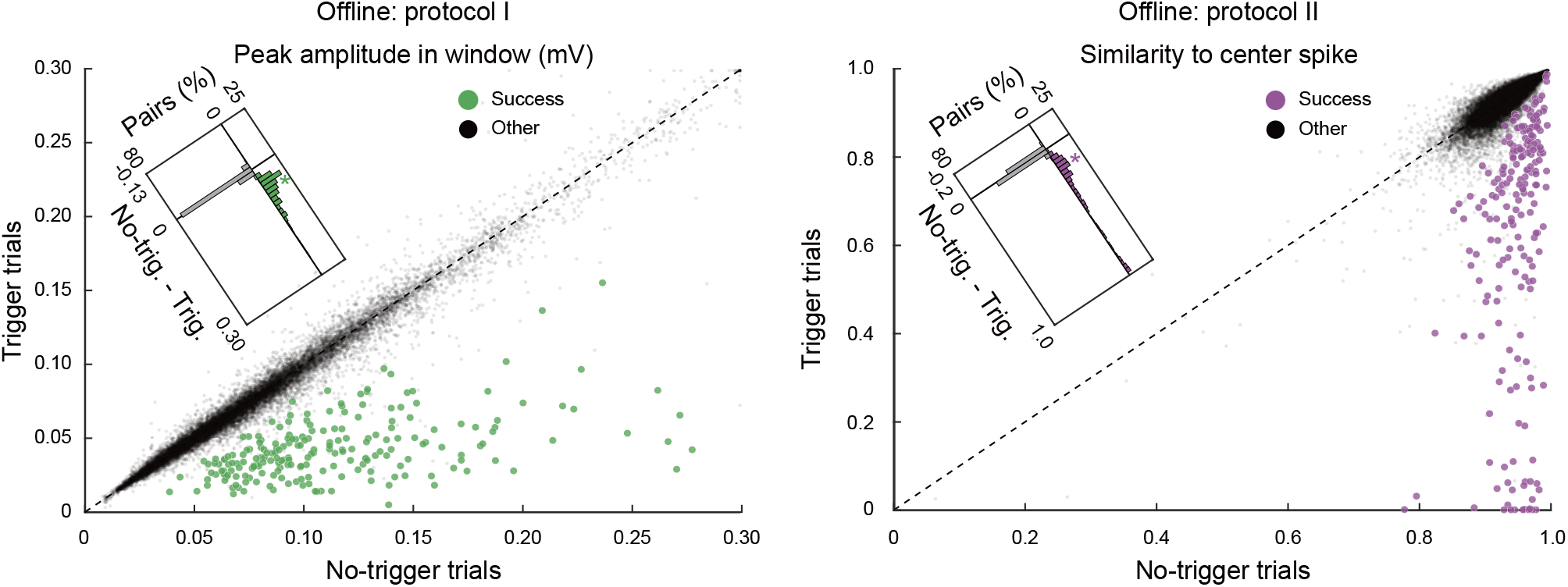
Confirmation of the collision judgement in the original scales. Our judgements of spike collision were based on the nonparametric statistics, AUC of ROC analysis between trigger and no-trigger trials. Here, we confirmed the segregation in the original scales of variables for judgements, the negative peak amplitude in protocol I, and the similarity to the center spike in protocol II. These variables in successfully identified pairs were confirmed to be smaller in trigger trials than in no-trigger trials (green and magenta dots below diagonal dashed lines; *p* = 4.8 × 10^−36^ in protocol I, *p* = 6.8 × 10^−35^ in protocol II, signed-rank test). The segregations in the successful pairs were larger than the others (*p* = 1.0 × 10^−135^ in protocol I, *p* = 1.2 × 10^−130^ in protocol II, rank-sum test for differences; diagonal insets).

**Fig. S6.**
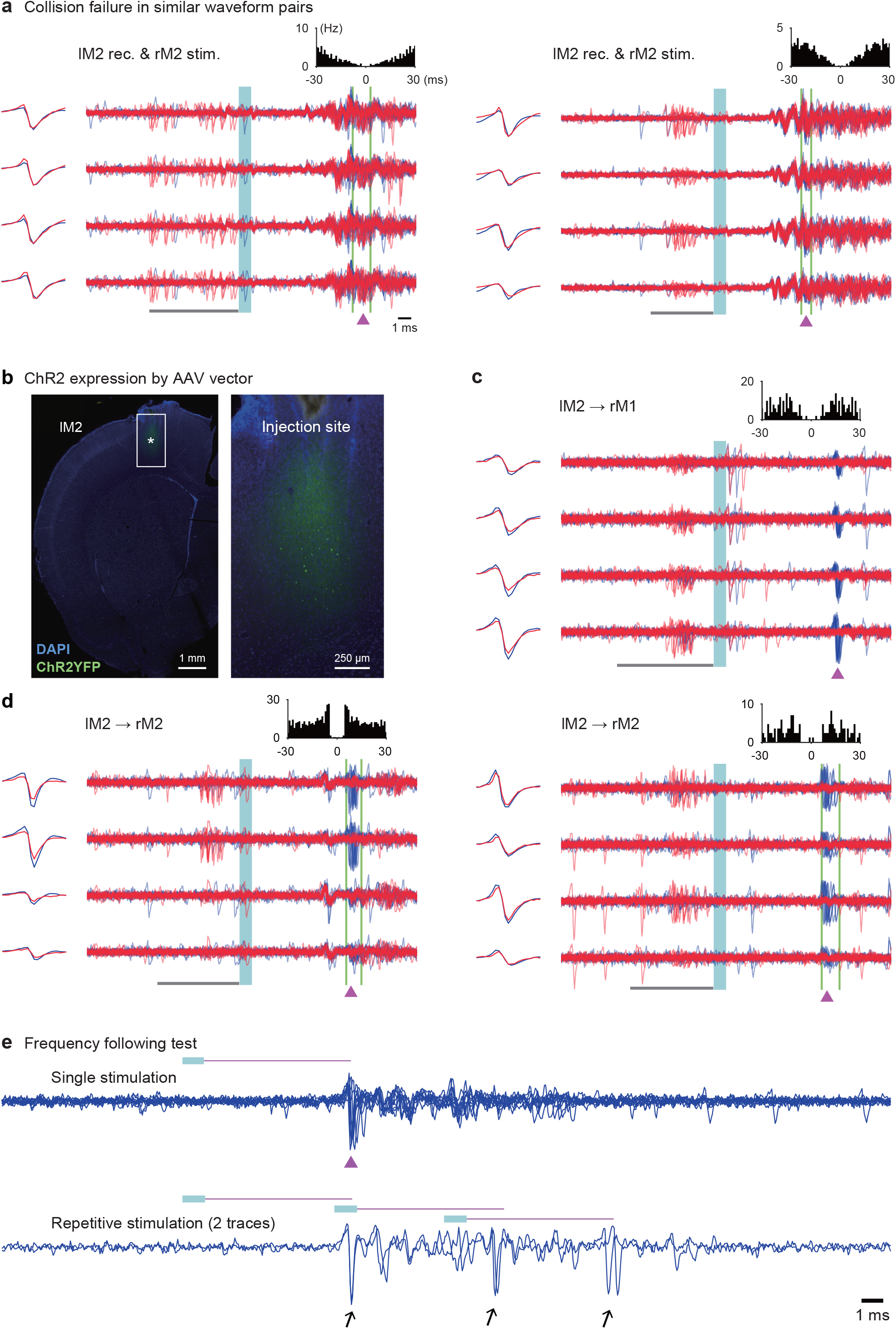
Other examples of spike collision tests using the *Multi-Linc* system. **a**, Two examples of neurons that did not pass the spike collision test despite their high similarity between trigger and evoked spikes. The waveforms of their trigger spikes were almost identical to those of evoked spikes (>0.9 in Pearson’s correlation coefficient), and the jitter of their evoked spikes satisfied the criterion (<0.25 ms). However, the AUC in amplitude/similarity between trigger and no-trigger trials did not satisfy the criterion (<5σ), rejecting their occurrence of spike collision. Data are shown in **Fig. 3a. b−d**, A preliminary example of a spike collision test in a wild-type (Long–Evans) rat with ChR2 expressed cortically via an AAV vector. **b**, Virally expressed ChR2-EYFP (green; immunostained) in the left M2 (left). An injection site (asterisk in a box) is shown magnified on the right. The *Multi-Linc* experiment was performed six weeks after cortical injection of AAVDJ-Syn-hChR2-EYFP (1.8 × 10^10^ vg/μl, 1 μl/site, provided by Dr. Kenta Kobayashi at the National Institute for Physiological Sciences, Japan) into the left M1 and M2. **c** and **d**, Successful spike collision tests using the *Multi-Linc* system in a session. Data are shown in **Fig. 3a. e**, Demonstration of a frequency-following test. The *Multi-Linc* system was configured to automatically perform several trials of the frequency-following test for evoked spikes if necessary. Upper, evoked spikes in response to single-pulse stimulation (blue) in a neuron that passed the spike collision test. A magenta triangle with a horizontal line indicates the timing of the center spike and its latency from the stimulation. Lower, similar spikes evoked exactly in response to each pulse of repetitive stimulation (arrows). It satisfied the frequency-following test.

**Table S1.** Summary of the identified projection neurons. Online: protocol I ∪ II

| Recording site | Stimulation site | Tested probe fiber | Number of projections | Latency (ms) | Jitter (ms) |
| --- | --- | --- | --- | --- | --- |
| IM1 | IM1 | 30 | 1 | 7.3 | 0.11 |
|  | IM2 | 6 | 0 |  |  |
|  | rM1 | 168 | 38 (33, 28) | 7.9 ± 1.5 | 0.11 ± 0.04 |
|  | rM2 | 104 | 7 (6, 6) | 7.4 ± 2.9 | 0.08 ± 0.06 |
| IM2 | IM1 | 38 | 7 (7, 5) | 3.9 ± 2.2 | 0.06 ± 0.05 |
|  | IM2 | 34 | 28 (21, 20) | 5.1 ± 2.0 | 0.07 ± 0.05 |
|  | rM1 | 162 | 22 (18, 19) | 8.9 ± 2.1 | 0.08 ± 0.03 |
|  | rM2 | 210 | 142 (113, 117) | 9.9 ± 2.4 | 0.11 ± 0.04 |
| rM1 | IM1 | 18 | 1 | 19.8 | 0.15 |
|  | IM2 | 2 | 0 |  |  |
|  | rM1 | 18 | 1 | 6.6 | 0.17 |
|  | rM2 | 2 | 0 |  |  |
| rM2 | IM1 | 2 | 0 |  |  |
|  | IM2 | 30 | 4 (3, 3) | 10.8 ± 0.7 | 0.18 ± 0.03 |
|  | rM1 | 2 | 0 |  |  |
|  | rM2 | 34 | 5 (5, 2) | 4.5 ± 1.5 | 0.09 ± 0.05 |

For each pair of a recording site and a stimulation site, we summarized the number of pairs of tested probe and stimulation, the number of identified projection neurons, the latency of evoked spikes (median ± quartile deviation), and the jitter of those (median ± quartile deviation) for online protocols I and II. The numbers in parentheses in "Number of projections" indicate those for offline judgements by protocols I and II, respectively. The latency and jitter were shorter for ipsilateral projections than for contralateral ones ( $p = 3.3 \times 10^{-9}$ and $p = 0.0081$ , respectively, rank-sum test).

**Table S2.** Summary of the statistical parameters.

| Figure | Test | Protocol | Descriptive statistics | DF | Test statistic | Effect size |
| --- | --- | --- | --- | --- | --- | --- |
| Fig. 3c | $\chi^2$ test | I | OR = 0.11 (0.08, 0.16) | 1 | $\chi^2 = 252.76$ | $\phi = 0.07$ |
| | | II | OR = 0.19 (0.13, 0.29) | 1 | $\chi^2 = 76.82$ | $\phi = 0.04$ |
| Fig. 3d | Pearson's correlation | I | $r = 0.57$ (0.46, 0.65) | 207 | Same as left | n/a |
| | | II | $r = 0.54$ (0.43, 0.63) | 200 | | |
| Fig. 4a | $\chi^2$ test | I | OR = 0.12 (0.04, 0.32) | 1 | $\chi^2 = 25.04$ | $\phi = 0.04$ |
| | | II | OR = 0.03 (0.00, 0.21) | 1 | $\chi^2 = 31.78$ | $\phi = 0.05$ |
| Fig. 4b | Rank-sum test<br>(waveform stability) | I | Success: M = 0.86 (0.84, 0.87)<br>Other: M = 0.74 (0.74, 0.75) | 5062 | $z = 13.04$ | $\delta = 0.53$ |
| | | II | Success: M = 0.88 (0.86, 0.89)<br>Other: M = 0.69 (0.68, 0.69) | 7999 | $z = 17.67$ | $\delta = 0.73$ |
| | Rank-sum test<br>(bias of peak-pattern) | I | Success: M = 0.37 (0.34, 0.41)<br>Other: M = 0.22 (0.21, 0.22) | 5062 | $z = 10.98$ | $\delta = 0.45$ |
| | | II | Success: M = 0.39 (0.36, 0.43)<br>Other: M = 0.13 (0.12, 0.13) | 7999 | $z = 17.29$ | $\delta = 0.71$ |
| Fig. 5a | Rank-sum test<br>(similarity of waveform) | I | Success: M = 0.97 (0.97, 0.98)<br>Other: M = 0.77 (0.77, 0.78) | 19511 | $z = 22.70$ | $\delta = 0.91$ |
| | | II | Success: M = 0.97 (0.97, 0.98)<br>Other: M = 0.78 (0.78, 0.78) | 34799 | $z = 21.66$ | $\delta = 0.88$ |
| Fig. 5b | Linear regression | I | $a_0 = 0.004$ (-0.003, 0.011) | 207 | $t = 1.05$ | $R^2 = 0.77$ |
| | | | $a_1 = 0.74$ (0.68, 0.79) | 207 | $t = 26.53$ | |
| | | II | $a_0 = 0.011$ (0.004, 0.018) | 200 | $t = 3.10$ | $R^2 = 0.77$ |
| | | | $a_1 = 0.69$ (0.63, 0.74) | 200 | $t = 25.65$ | |
| | Signed-rank test<br>(evoked vs. trigger) | I | M = 0.03 (0.02, 0.03) | 208 | $z = 11.93$ | $r = 0.83$ |
| | | II | M = 0.02 (0.02, 0.02) | 201 | $z = 11.71$ | $r = 0.82$ |
| Fig. S5 | Signed-rank test<br>(no-trigger vs. trigger) | I | M = 0.06 (0.05, 0.07) | 208 | $z = 12.53$ | $r = 0.87$ |
| | | II | M = 0.21 (0.19, 0.25) | 201 | $z = 12.32$ | $r = 0.87$ |
| | Rank-sum test | I | Success: M = 0.059 (0.053, 0.067)<br>Other: M = $1.0 \times 10^{-4}$<br>($5.0 \times 10^{-5}$, $1.4 \times 10^{-4}$) | 19511 | $z = 24.80$ | $\delta = 1.00$ |
| | | II | Success: M = 0.21 (0.19, 0.25)<br>Other: M = $6.6 \times 10^{-4}$<br>($5.6 \times 10^{-4}$, $7.7 \times 10^{-4}$) | 34799 | $z = 24.32$ | $\delta = 0.99$ |

For each figure, we summarized the type of test, offline protocol, descriptive statistics, degree of freedom (DF), test statistic, and effect size. All tests were two-tailed. For descriptive statistics, OR, M, and $a_k$ are odds ratio, median, and k-th regression coefficient, respectively. The numbers in parentheses in "Descriptive statistics" indicate 95% confidence intervals calculated by bootstrap. Their p-values are given in the main text or this supplementary information.

